# A Genome-Scale TF-DNA Interaction Network of Transcriptional Regulation of Arabidopsis Primary and Specialized Metabolism

**DOI:** 10.1101/2021.05.13.443927

**Authors:** Michelle Tang, Baohua Li, Xue Zhou, Tayah Bolt, Jia Jie Li, Neiman Cruz, Allison Gaudinier, Richard Ngo, Caitlin Clark-Wiest, Daniel J. Kliebenstein, Siobhan M. Brady

## Abstract

In single-celled microbes, transcriptional regulation by single transcription factors is sufficient to shift primary metabolism. Corresponding genome-level transcriptional regulatory maps of metabolism reveal the underlying design principles responsible for these shifts as a model in which master regulators largely coordinate specific metabolic pathways. Relative to individual microbes, plant metabolism is more complex. Primary and specialized metabolism occur within innumerable cell types, and their reactions shift depending on internal and external cues. Given the importance of plants and their metabolites in providing humanity with food, fiber and medicine, we set out to develop a genome-scale transcriptional regulatory map of Arabidopsis metabolic genes. A comprehensive set of protein-DNA interactions between *Arabidopsis thaliana* transcription factors and promoters of primary metabolism and specialized metabolism were mapped. To demonstrate the utility of this resource, we identified and functionally validated regulators of the TCA cycle. The resulting network suggests that plant metabolic design principles are distinct from that of microbes. Instead, metabolism appears to be transcriptionally coordinated via developmental- and stress-conditional processes that can coordinate across primary and specialized metabolism. These data represent the most comprehensive resource of interactions between TFs and metabolic genes in plants.

## INTRODUCTION

Metabolism is the fundamental biological process underpinning all cellular functions. An organism’s metabolism comprises individual biochemical reactions organized into metabolic pathways where metabolites are sequentially transformed in increasing or decreasing complexity by enzymes. In conjunction, the primary metabolic pathways create the cellular building blocks and directly contribute to the inter-conversion of chemicals into energy currency. In plants, and most other organisms, primary metabolites serve as precursors to secondary, or specialized, metabolites crucial to the organism’s interaction with its environment. Plant specialized metabolites serve many functions, including defending plants from predators and pathogens, attracting symbiotic organisms and promoting interactions with pollinators.

To properly function, metabolic pathways must be intricately orchestrated to maintain the homeostasis necessary for growth and are in turn dependent on the organism’s developmental stage and environment. Thus, it is critical to understand how metabolic pathways are regulated to maximize our ability to predict and manipulate an organism’s genotype to phenotype matrix.

Metabolism is known to be regulated by mechanisms that span the central dogma from mRNA transcription to protein post-translational modifications with the best-studied regulatory mechanisms being post-translational modification and allosteric feedback of enzymes (Nielsen, 2017). Adding to this understanding, systems biology and genetic approaches in single-celled organisms demonstrate the importance of transcriptional regulation. In *Saccharomyces cerevisiae* and *Escherichia coli*, these systems approaches integrate chromatin immunoprecipitation, transcriptomic experiments and *in silico* models to elucidate transcription factor (TF) – enzyme promoter regulatory interactions and ultimately, investigation of genome-scale regulatory networks of global metabolism (Barrett et al., 2005; Fang et al., 2017; Ihmels et al., 2002; 2004; Lempp et al., 2019). Studies in these organisms have resulted in a model where metabolic networks are organized into distinct transcriptional modules that control specific cellular processes (Ihmels et al., 2002; 2004). Whether the same principles apply in multicellular organisms remains to be determined.

Relative to *S. cerevisiae* and *E. coli*, the genomes of multicellular organisms encode many more genes including TFs, enzymes and in plants, relative to animals, a further expansion of both enzyme and transcription factor families linked to metabolism. The acquisition of multicellularity also enables partitioning of function across cell types. Thus, multicellular organisms likely have more complex transcriptional and metabolic regulation in comparison with single cell microbes. However, few studies exist which systematically characterize the complexity of metabolic networks from the perspective of transcriptional regulation in multicellular organisms. Even fewer studies have explored the regulatory interconnection between regulation of central and specialized metabolism in multicellular organisms, where specialized metabolism is a critical component of the organism’s response to the environment. Instead, the majority of studies on plant metabolism have focused on transcriptional regulation of individual pathways (Bonawitz et al., 2012; Dolan et al., 2017; Gaudinier et al., 2018; Kim et al., 2015; Li et al., 2014; 2018).

Recent work is beginning to show that plant primary and specialized metabolism are highly interconnected. The potential for interconnected regulation of central carbon and specialized metabolism is revealed by transcriptional profiling of mutants in TFs, MYB28 and MYB29 that regulate GSL biosynthesis, a specialized metabolic pathway. This work showed that, as expected, these TFs regulate the glucosinolate biosynthetic pathway, but also affect several primary metabolic pathways which synthesize precursors of glucosinolates, including methionine biosynthesis, tryptophan biosynthesis, the TCA cycle, sulfur metabolism and folate metabolism (Malitsky et al., 2008; Sønderby et al., 2010). Another example of TFs that affect both central carbon and specialized metabolism are the Mediator complex, a multi-subunit transcriptional co-regulator that interacts with other transcription factors and the general transcription machinery to regulate transcription (Tsai et al., 2014). In Arabidopsis, different subunits of the Mediator complex are involved in fatty acid biosynthesis (Kim et al., 2016) and the synthesis of specialized metabolites known as phenylpropanoids (Bonawitz et al., 2012; Dolan and Chapple, 2018; Dolan et al., 2017; Stout et al., 2008). These observations suggest that transcriptional regulation of plant metabolism may be highly coordinated and differ from the pathway-specific transcriptional regulatory modules as observed in yeast and *E. coli*. Thus, we hypothesized that Arabidopsis metabolism is orchestrated by TFs that regulate both primary and specialized metabolism.

Testing this hypothesis requires a large-scale dataset linking as many TFs to primary metabolism promoters as possible. Presently, large-scale regulatory transcriptional regulatory networks of plant metabolism can be inferred computationally with tools and publicly available datasets including RNA-Seq, ChIP-Seq, DAP-Seq and protein binding microarrays (Kulkarni et al., 2018; O’Malley et al., 2016; Weirauch et al., 2014). However, these datasets are limited to a subset of TFs, with the Arabidopsis cistrome dataset consisting of 529 Arabidopsis TFs (O’Malley et al., 2016). We complement these approaches using enhanced yeast one-hybrid assays (Gaudinier et al., 2011) to systematically screen for interactions between approximately 85% of all characterized and putative TFs in Arabidopsis and 226 promoters of enzyme genes in 11 central carbon metabolic pathways and of a specialized metabolic pathway, glucosinolate biosynthesis. By developing a comprehensive dataset linking transcription factors to metabolic genes, we can begin addressing key questions in transcriptional regulation of multicellular metabolism: are metabolic pathways linked by common precursors coordinately regulated by single TFs? Do TFs confer environmental conditionality of primary metabolism as occurs for specialized metabolism?

This dataset showed that 90% of TFs bind to promoters of metabolic genes with all TFs binding to promoters of multiple metabolic pathways. Co-expression analyses support a regulatory model whereby the majority of TFs influence more than one metabolic pathway. We validated this hypothesis and the dataset by expression profiling inducible lines of four TFs that bind to promoters from a range of metabolic pathways. This showed that the predicted target enzyme genes in the central carbon metabolic pathways were enriched among the differentially expressed genes of the TFs we examined. Additionally, we explored combinatorial regulation of a single pathway, the tricarboxylic acid (TCA) cycle, given its interconnection with other central metabolic pathways. This dataset provides a unique resource to extend our understanding of how metabolism is regulated in multi-cellular organisms.

## RESULTS

### Genome-scale identification of TFs involved in primary and secondary metabolism

To identify transcription factors that potentially regulate Arabidopsis primary metabolism, we conducted enhanced yeast one-hybrid (Y1H) screens (Gaudinier et al., 2011) with 224 promoters of enzyme-encoding genes involved in the TCA cycle, glycolysis and gluconeogenesis, pentose phosphate pathway, glutamine synthetase/glutamine oxogluatarate aminotransferase (GS-GOGAT) cycle, shikimate pathway, and most amino acid biosynthesis pathways with the exception of the aromatic amino acids (Figure 1, Table S1). We refer to this collective group of promoters as central carbon promoters (CCP). We also included promoters of genes involved in aliphatic GSL biosynthesis (Li et al., 2014), to investigate transcriptional coordination of primary and secondary metabolism. To enable genome-level detection of TFs involved in primary metabolism, we extended our original collection of 812 TFs by cloning an additional 1227 TFs into activation domain fusion vectors compatible with our enhanced yeast one-hybrid system (Table S2) (Gaudinier et al 2011, Li et al. 2014, Gaudinier et al 2018). The final Arabidopsis TF collection contains 2039 TFs and represents over 80% of all characterized and putative TFs in Arabidopsis (Pruneda Paz et al 2015; see https://genomecenter.ucdavis.edu/yeast-one-hybrid-services).

**Figure 1.**
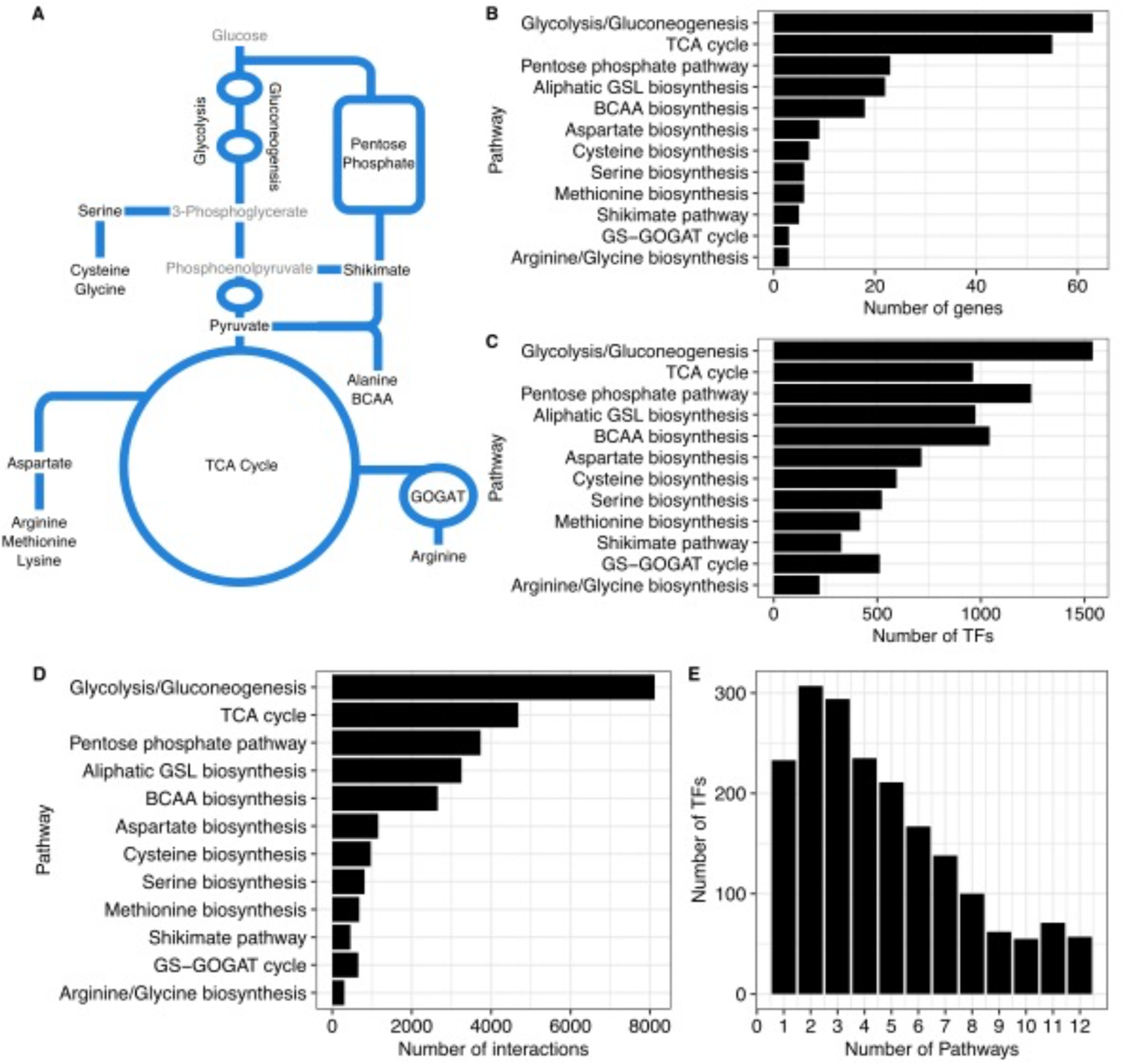
Summary of TF-promoter interactions of central carbon and specialized metabolism. (A) The simplified biochemical network comprises amino acid biosynthetic and respiratory pathways in central carbon metabolism and a specialized metabolic pathway, aliphatic glucosinolate biosynthesis, studied in this paper. (B) Each pathway in central carbon and specialized metabolism vary in gene number. (C and D) Y1H identified TFs that interact with promoters of each pathway in central carbon and in the specialized metabolic pathway. (E) Majority of TFs bind to promoters from two or more metabolic pathways.

**Table 1.** Associated metabolic pathways of promoters bound by transcription factors tested in conditional GR-induction assays.

|  | Aliphatic GSL<br>biosynthesis<br>(22) | Arginine/Glycine<br>biosynthesis<br>(3) | Aspartate<br>biosynthesis<br>(9) | BCAA<br>biosynthesis<br>(18) | Cysteine<br>biosynthesis<br>(8) | Glycolysis/<br>Gluconeogenesis<br>(64) | GS-GOGAT<br>cycle<br>(3) | Methionine<br>biosynthesis<br>(6) | Pentose<br>phosphate<br>pathway<br>(24) | Serine<br>biosynthesis<br>(6) | Shikimate<br>pathway<br>(5) | TCA cycle<br>(56) |
| --- | --- | --- | --- | --- | --- | --- | --- | --- | --- | --- | --- | --- |
| CHA19 | 9 | 1 | 2 | 6 | 1 | 19 | 2 | 1 | 8 | 3 | 1 | 15 |
| ENAP1 | 18 | 3 | 4 | 8 | 6 | 49 | 2 | 3 | 21 | 4 | 1 | 33 |
| LBD16 | 9 | 2 | 5 | 15 | 6 | 41 | 1 | 5 | 20 | 4 | 2 | 24 |
| WRI3 | 0 | 0 | 0 | 0 | 0 | 8 | 0 | 0 | 3 | 0 | 0 | 7 |

Summary of the number of targets in each metabolic pathway of CHA19, ENAP1, LBD16 and WRI3 from Y1H assays. The number of promoters of each metabolic pathway assayed are in parentheses.

From the binding data obtained via the genome-scale Y1H, we queried how promoters and metabolic pathways are organized. We detected 27,485 interactions between 1930 TFs and 220 promoters across the 12 pathways surveyed (Figure 1A-C, Figure S1, Table S3). Per promoter, we identified 1-509 TF interactions, with an average of 125 TFs binding to a promoter. To visualize if most TFs are specific to individual pathways or connect to multiple metabolic pathways, we mapped TF-promoter interactions relative to their respective pathways (Figure 1B and 1C). While we observed a general positive relationship between the number of interactions and the number of promoters screened for each pathway (Figure 1B), the number of TFs identified did not scale with the number of promoters in each pathway. For instance, we detected approximately the same number of TFs binding to promoters associated with serine biosynthesis and the GOGAT cycle even though there are twice as many genes encoding enzymes in serine biosynthesis as in the GOGAT cycle. While the large number of TFs found to bind to metabolic promoters could suggest that specific groups of TFs target unique pathways, the majority of TFs (∼88%) in our network bind to promoters of two or more pathways (Figure 1C). Indeed, only 10% of the TFs bound to promoters of a single pathway. This pattern of binding suggests TFs may regulate multiple pathways to coordinate metabolic control (Figure 1C). We next queried the percentage of TFs that are shared between metabolic pathways. In all pairwise combinations, the number of TFs common between any two pathways is significantly greater than expected (Fisher’s exact test, FDR < 0.0001), signifying that transcriptionally-mediated interconnection may be an emergent property of plant metabolism (Figure 2, Table S4). Such coordination may be guided to control a particular biological process or molecular function that requires a subset of metabolic pathways. However, Gene Ontology (GO) enrichment analysis for each pathway did not provide any clarity, likely due the fact that most TFs do not have known functions or metabolism-associated annotations (Table S5).

**Figure 2.**
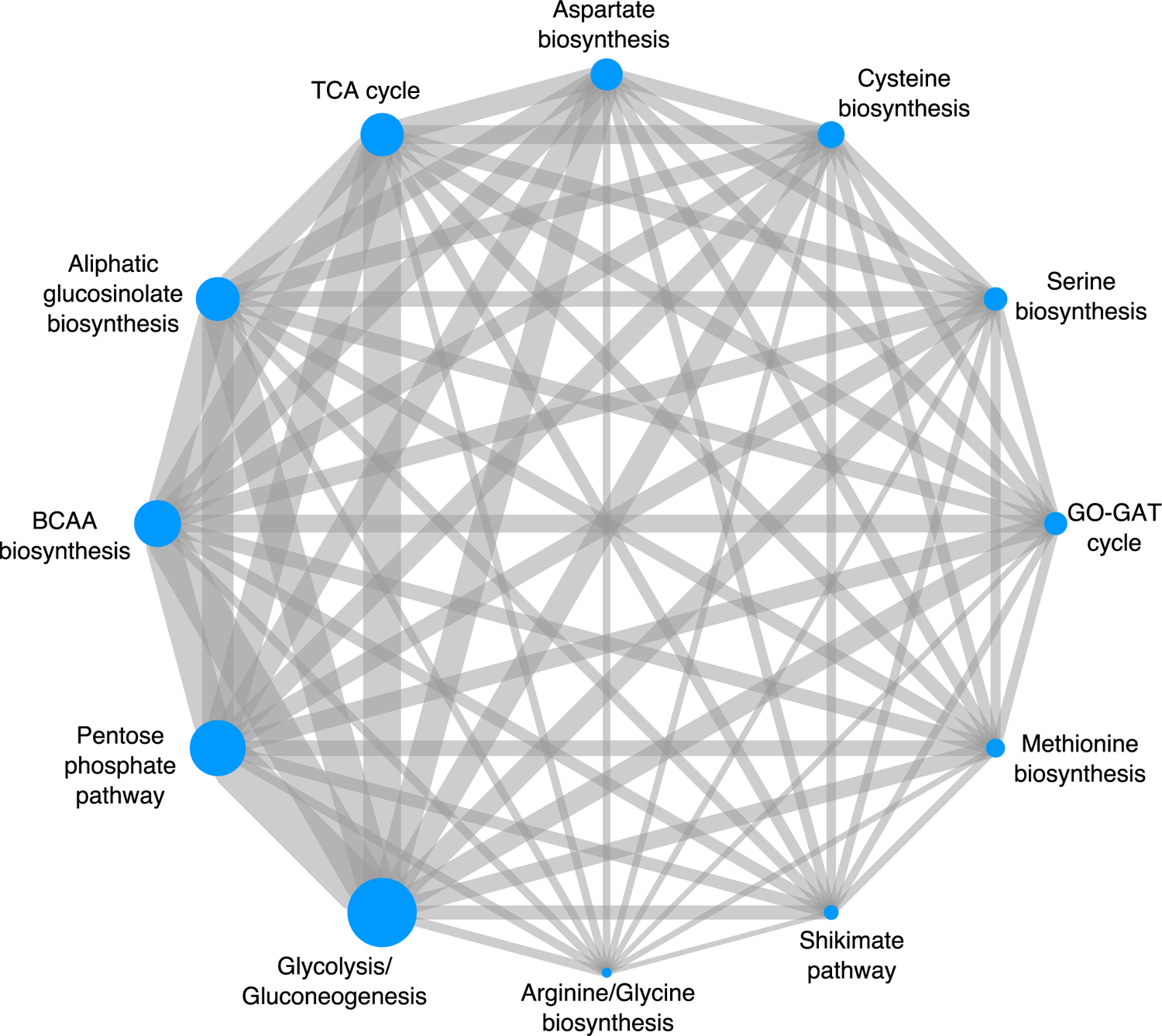
Pairwise association of TFs between metabolic pathways. The number of TFs shared between metabolic pathways is greater than expected by chance for all combination of pairs of metabolic pathways (Table S4). The size of the nodes corresponds to the number of TFs identified for each pathway. The width of the edge linking two metabolic pathway indicates the number of TFs shared between the two pathways.

We next examined how our representative specialized metabolic pathway, aliphatic GSL biosynthesis, is integrated with central carbon metabolism within the TF-DNA interaction network. Among the 974 TFs that bind to aliphatic GSL gene promoters, 933 of those TFs (∼95%) also bind to promoters from a central carbon pathway. In our pairwise comparison of pathways based on TF binding, the central carbon metabolic pathways that shared the most TFs with aliphatic GSL based on the level of enrichment using a hypergeometric test are the pathways that synthesize the sulfur containing precursors needed to synthesize aliphatic GSL (methionine and cysteine). This signifies that although the number of TFs shared between pathways are higher than expected across all pairwise combination of pathways, there is still regional enrichment wherein biosynthetic pathways that are closest within the metabolic network share more TFs than biosynthetic pathways more distant in the metabolic network.

### Coordinate transcriptional regulation of central carbon metabolism

This global analysis of TF-metabolic promoter interactions suggests coordinate transcriptional regulation of central carbon and specialized metabolic pathways. However, genome-scale assays like the yeast-one hybrid suffer from both false positive and false negatives. We therefore tested the regulatory capacity of four transcription factors that bind to gene promoters from multiple metabolic pathways: *CHROMATIN REMODELING 19 (CHA19)*, *EIN2 NUCLEAR ASSOCIATED PROTEIN 1 (ENAP1)*, *LATERAL ORGAN BOUNDARIES-DOMAIN 16 (LBD16)* and *WRINKLED 3 (WRI3)* (Table 1). WRI3 binds to promoters in the TCA cycle, glycolysis/gluconeogenesis and pentose phosphate pathway while CHA19, ENAP1 and LBD16 bind to promoters from all 12 pathways at varying degrees (Table 1, Figure 3A). We reasoned that conditional overexpression of these transcription factors would reveal their sufficiency to regulate gene expression in these pathways. Additionally, we utilized the fact that dark grown plant seedlings are heterotrophic and do not shift to autotrophy (photosynthesis) until exposure to light. Thus, we tested the regulatory capacity for these TFs in dark-grown seedlings. We reasoned that carbon metabolism relating to photosynthesis and carbon fixation would be suppressed in dark-grown seedlings, and seedlings would maintain heterotrophy during catabolism of maternal stores to provide energy for seed germination and growth. *CHA19*, *ENAP1*, *LBD16* and *WRI3* coding regions were fused to a dexamethasone (Dex)-controlled glucocorticoid receptor. Gene expression was profiled by RNA-sequencing 24 hours after mock treatment or 10 uM dexamethasone induction in dark-grown six-day old Arabidopsis seedlings. Hundreds to thousands of genes were differentially expressed in response to transcription factor induction across multiple insertion lines (Figure 3B, Table S6). There was a significant enrichment of Y1H network gene targets amongst these <u>d</u>ifferentially <u>e</u>xpressed <u>g</u>enes (DEGs) for *GR-ENAP1; GR-LBD16* and *GR-WRI3* (one-tailed Fisher’s exact test, P < 0.05).

**Figure 3.**
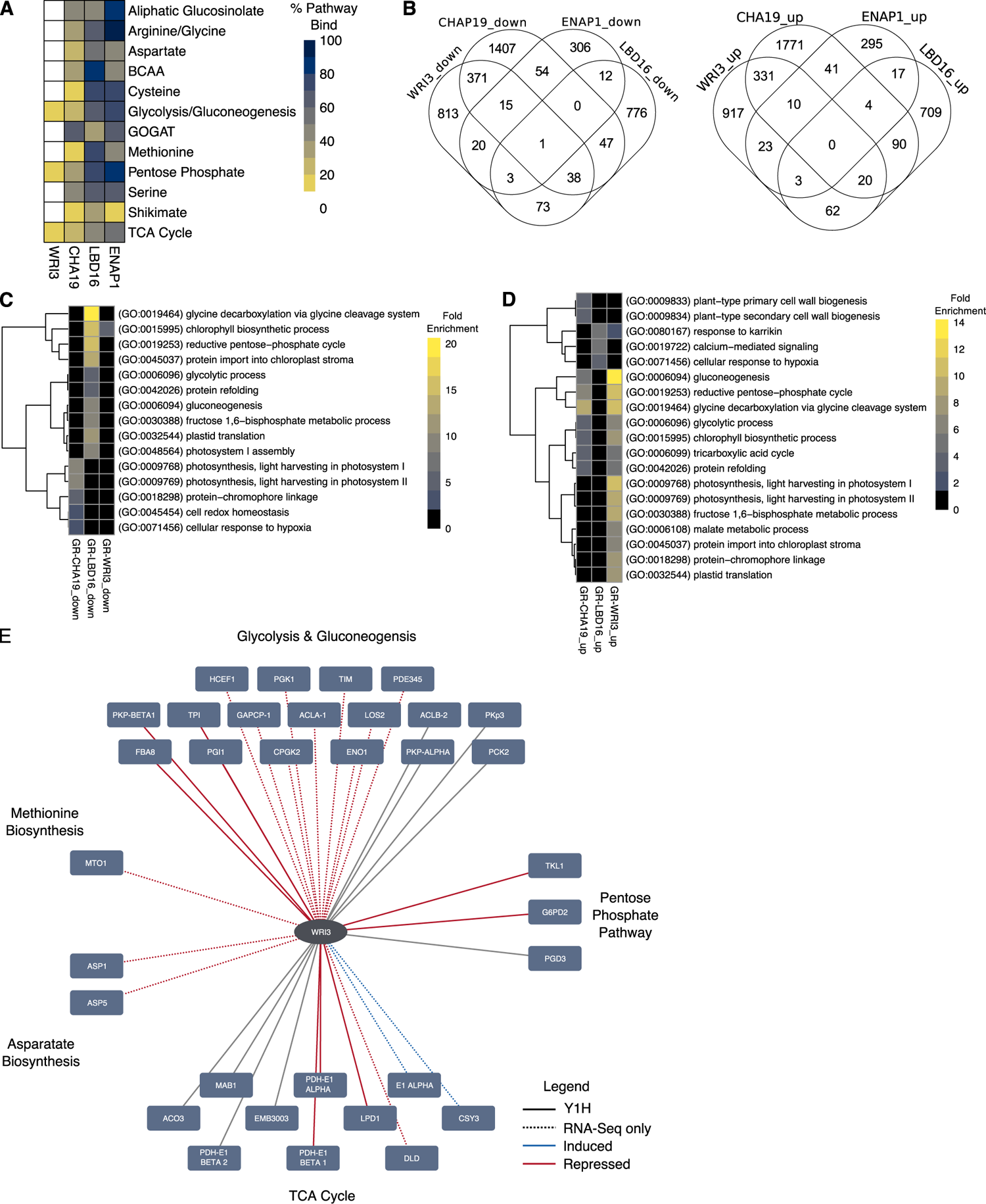
Validation of regulatory interactions via transcriptomics of GR-TFs. (A) CHA19, ENAP1, LBD16 and WRI3 vary in the pathways and the number of genes targeted based on Y1H. (B) Thousands of genes were differentially expressed in Dex-induced GR-TFs compared to GR-TFs under mock conditions. (C) Gene ontologies enriched in the up-regulated DEGs included metabolic pathways in central carbon metabolism. (D) Gene ontologies enriched in the down-regulated DEGs related to central carbon metabolism were found only in GR-LBD16. (E) Among the genes in the Y1H network, glycolysis/gluconeogenesis and TCA cycle genes were enriched in DEGs of GR-WRI3.

GO and pathway enrichment analysis further corroborated the effect of these TFs on central carbon metabolism. Gene ontologies associated with the reductive pentose phosphate pathway, glycine biosynthesis, and cysteine biosynthesis were over-represented more than expected by chance in the significantly up-regulated DEGs upon GR induction of *CHA19, LBD16* and *WRI3* (Figure 3C). Only *GR-LBD16* had down-regulated DEGs enriched for GO terms involving CCP pathways (glycine, reductive pentose phosphate pathway, glycolysis and gluconeogenesis) (Figure 3D). Pathway enrichment analysis (Methods) revealed in particular, significant enrichment of DEG in the TCA cycle for *GR-WRI3*. 71.4% of the Y1H-predicted *WRI3* TCA cycle targets were recovered in the RNA-Seq analysis (Figure 3E, P=0.0006, Fisher’s exact test). Additionally, we found other TCA cycle genes, totaling 28.5% of TCA cycle genes, to be enriched in the set of significant Dex-dependent genes (Figure 3E, P = 0.0018, Fisher’s exact test). These data collectively demonstrate that our TF-DNA interaction network was able to predict *in planta* regulatory interactions, in line with mapped networks for other biological processes (Gaudinier et al., 2018; Li et al., 2014; Taylor-Teeples et al., 2014; Truskina et al., 2021).

In most heterotrophic and autotrophic eukaryotes, the TCA cycle occurs predominantly in mitochondria. Plants, however, have expanded this repertoire such that TCA cycle isoforms also function in the cytosol, peroxisomes and plastids, thus allowing pathway interactions with photosynthesis, photorespiration and nitrogen assimilation. These isoforms create the potential for localized bypasses, resulting in cyclic and non-cyclic fluxes in the TCA cycle to optimize metabolism (Araújo et al., 2012; Tcherkez et al., 2009). Given the relatively large number of TCA cycle genes differentially expressed upon GR-induction of *WRI3*, we inspected their subcellular localization. The TCA cycle DEGs of GR-WRI3 are specifically localized in the mitochondria and plastids (Figure 4A). This showed that only the mitochondrial and plastidic TCA pathways were being influenced by WRI3 indicating the potential for TFs to differentially modulate pathways targeted to different subcellular compartments (Figure 4A, P=0.01159 (mitochondria) and P=0.001431 (plastid), respectively, Fisher’s exact test, Methods). Overall, the inducible constructs support the Y1H binding results and the hypothesis of coordinated regulation across metabolic pathways.

**Figure 4.**
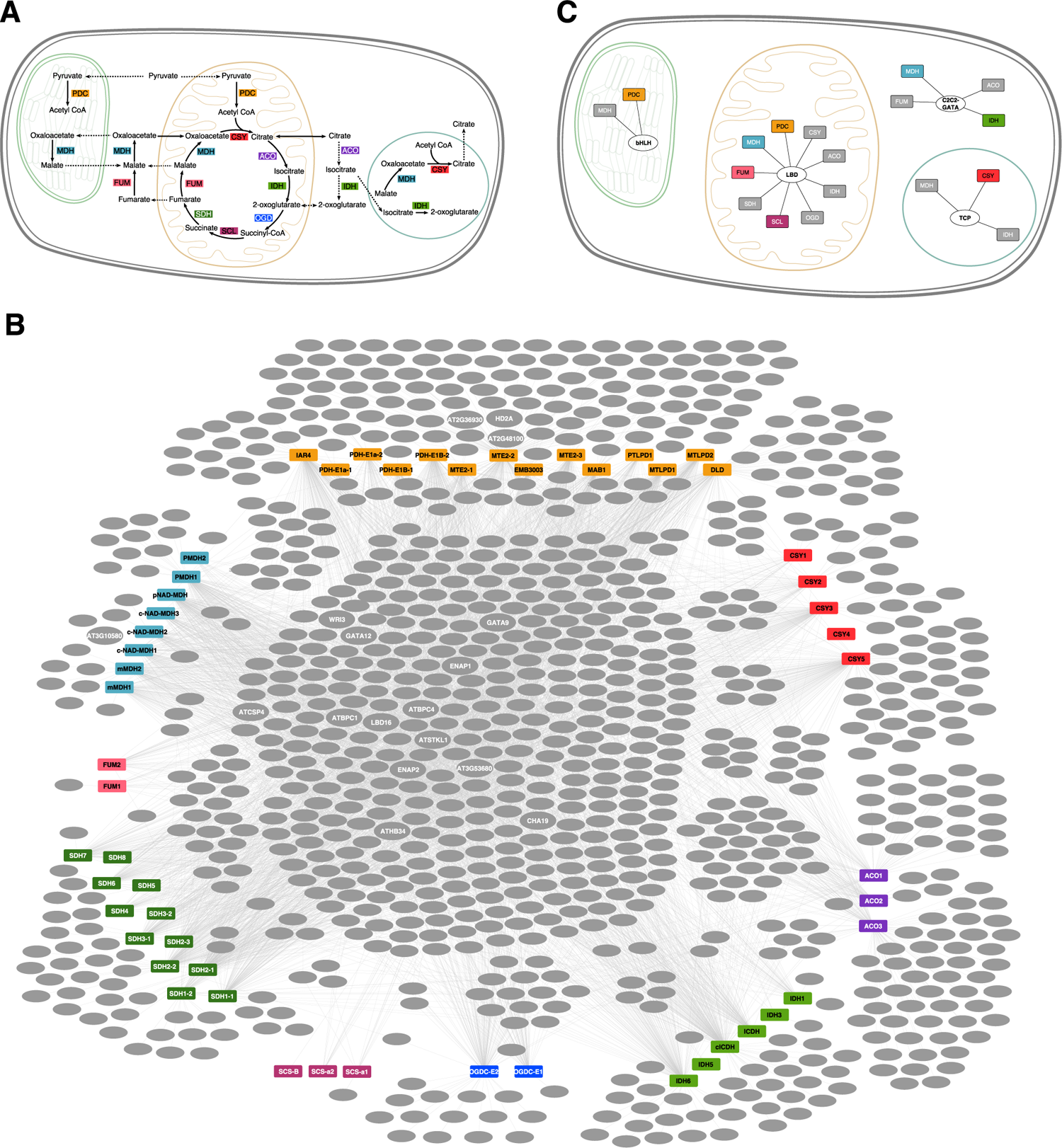
Arabidopsis TCA cycle and its Y1H network. (A) TCA cycle-associated metabolites and isozymes in the plastid (green), mitochondrion (orange), peroxisome (blue) and cytosol of a plant cell allow for non-cyclic flux, thus increasing the flexibility of the pathway. Organelles are not drawn to scale. (B) Y1H network shows the interactions between transcription factors and promoters of TCA cycle genes. Promoters are colored rectangles. The following colors correspond with the metabolic pathways: Orange PDC, pyruvate dehydrogenase; red CSY, citrate synthase; purple ACO, aconitase; light green IDH, isocitrate dehydrogenase; blue OGD, oxoglutarate dehydrogenase; light purple SCL, succinyl-CoA ligase; green SDH, succinate dehydrogenase; pink FUM, fumarase; and light blue MDH, malate dehydrogenase. Grey ovals denote transcription factor, and grey edges indicated interactions detected via Y1H. Tested TFs are labelled. See Figure S1 for full diagram. (C) TF family (oval) enrichment is modularly organized by the interaction of cellular localization and enzyme (rectangles). Colored enzymes indicate significant TF family enrichment in cellular compartment (adjusted P < 0.05, Fisher’s exact test).

### Uncovering regulators of the Arabidopsis TCA cycle

To develop a deeper understanding of a single metabolic pathway, we focused on the TCA cycle that converts nutrients into carbon skeletons for more complex biomolecules and reduces electron carriers for ATP synthesis. These functions are carried out by eight enzymes: citrate synthase, aconitase, isocitrate dehydrogenase, oxoglutarate dehydrogenase, succinyl-CoA ligase, succinate dehydrogenase, fumarase and malate dehydrogenase, many of whom have organelle-specific isoforms (Figure 4A). Pyruvate dehydrogenase serves as a critical link between glycolysis and the TCA cycle, which together, form the central hub of carbon metabolism. Thus, the plant TCA cycle comprises organellar-specific isoforms, connects with many metabolic processes, and its activity likely functions in different ways in cells, tissues, organs and the plant’s response to the environment. We therefore use the TCA cycle as a reference point to examine how transcription factors function to organize the TCA cycle and ultimately how this regulation coordinates plant growth, development and response to the environment.

We first mapped a TCA cycle sub-network consisting of 4,684 interactions between 962 TFs and 55 TCA cycle genes (Figure 4B, Figure S1, Table S1 and S3). Within this sub-network, we tested the hypothesis that certain TF families coordinate specific isoforms within specific cellular compartments. Enrichment was constrained to specific enzyme isoforms targeted to defined cellular compartments and not for all TCA enzymes in that compartment (Figure S2). For example, TFs in the LATERAL ORGAN BOUNDARIES DOMAIN (LBD) family were enriched for binding to promoters of mitochondrial-localized forms of pyruvate dehydrogenase, succinyl-CoA ligase, fumarase, and malate dehydrogenase (Figure 4C, Figure S2, Table S7). These data suggest that the transcriptional control of the TCA cycle is organized around the intersection of enzyme complex and sub-cellular compartment.

As metabolic priorities differ between plant cell types and shift while adapting to environmental changes, we predicted that a given TF’s regulation of TCA cycle genes will be structured by development and the environment. To test this prediction of context-dependency, we estimated the Pearson correlation coefficients of TFs and their TCA targets across five microarray datasets that surveyed different developmental processes and environments: (i) plant development (Schmid et al., 2005); (ii) root cell type development (Birnbaum et al., 2003; Brady et al., 2007; Lee et al., 2006; Levesque et al., 2006); (iii) pollen development (Honys and Twell, 2004; Qin et al., 2009); (iv) osmotic stress (Kilian et al., 2007); (v) salt stress (Kilian et al., 2007) (Tables S8, Key Resources). If our hypothesis of conditional regulation was true, we expected to detect few interactions in common across the five microarray datasets. 1,046 TF/TCA target interactions from the TCA cycle sub-network (∼26%) were highly correlated (|r| > 0.8) in one or more microarray datasets (Figure S3). Of these highly correlated TF/TCA target interactions, 797 interactions (∼76%) were exclusive to only one of the five microarray datasets. Roughly 23% (237) of the highly correlated interactions between TF and TCA targets were found in two microarray datasets with the vast majority of these being an overlap between related salt and osmotic stress microarrays. The absence of universal TF/TCA target interaction and the high percentage of interactions being specific to a developmental or stress dataset align with our hypothesis that regulation of TCA cycle genes is highly conditional.

### Seventeen Transcription Factors Contribute to TCA-Cycle Dependent Plant Growth

To test the function of transcription factors within this TCA cycle sub-network, we developed a system to test the phenotypic consequences of defects in the TCA cycle. Dark grown Arabidopsis seedlings utilizes requires the TCA cycle to catabolize seed stores and exogenous carbohydrates and lipids for growth (Angelovici et al., 2011; Lee et al., 2010; Padmasree et al., 2002; Zakhartsev et al., 2016). Wild-type Col-0 seedlings are etiolated with long hypocotyls and a short root in the dark (Figure 5A). Our previous transcriptome profiling experiment demonstrates that WRI3 is sufficient to regulate TCA cycle gene expression in the dark. In agreement with these data, the hypocotyl and root length of the *wri3* loss-of-function mutant allele is shorter compared to wild type (Figure 5C-E). This suggests that we can utilize hypocotyl and root length as a proxy for alterations in the respiratory/TCA cycle output in plant growth and development.

**Figure 5.**
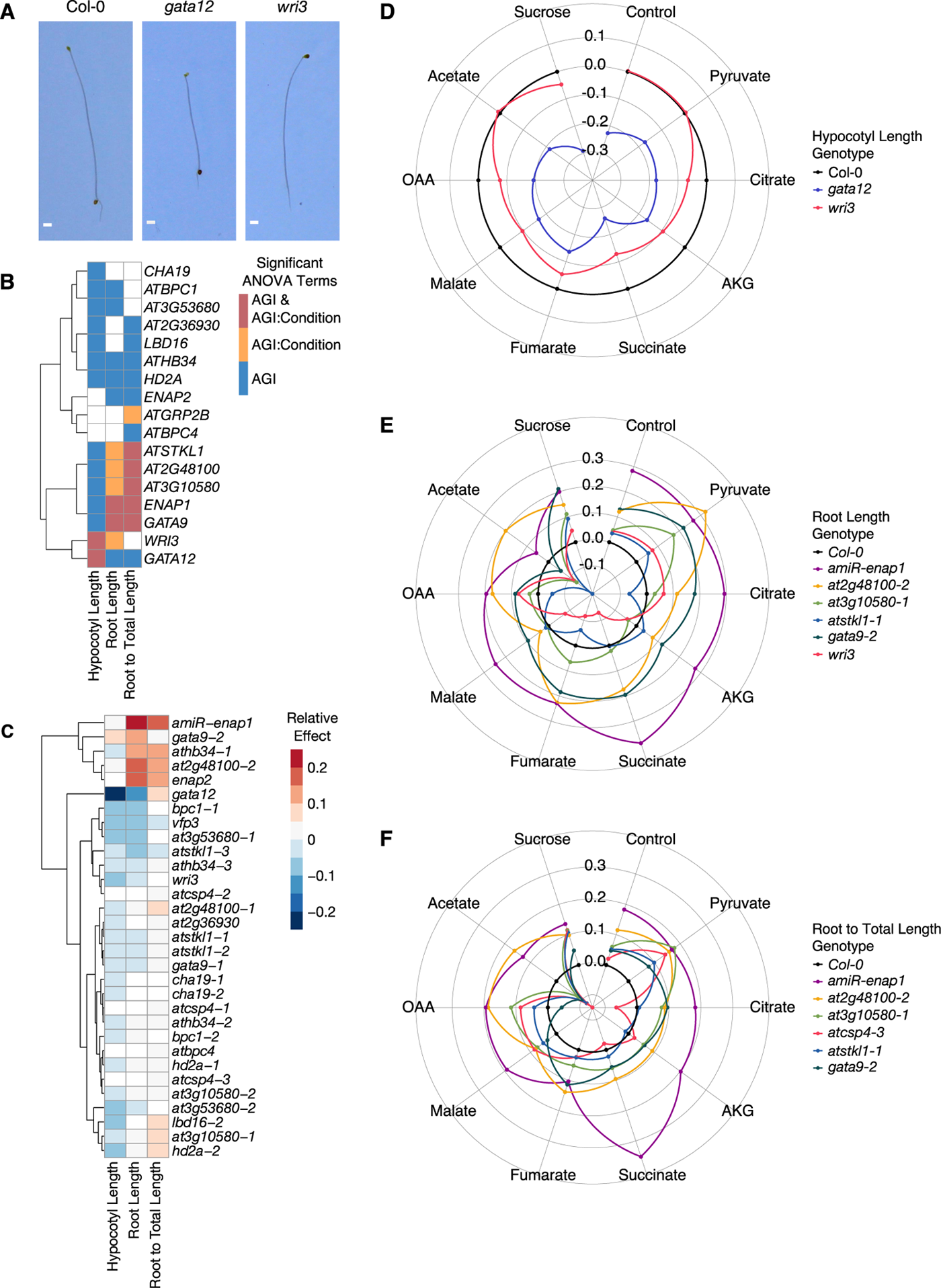
Characterization of TCA cycle function in transcription factor mutant alleles. (A) Representative 5-day-old seedlings of wild-type Arabidopsis thaliana Col-0 and TF mutants *gata12* and *wri3* grown in the dark. Scale bar, 1 mm. (B) Heat map indicates which TFs significantly affects hypocotyl length, root length and the composite trait of root to total length in dark-grown seedlings and whether the effects of TFs are dependent on condition. TFs are hierarchically clustered using Euclidean distance. (C) Heat map of the average relative effects of TF mutant alleles on hypocotyl length, root length and the ratio of root to total length reveal that TF lesions significantly perturbed TCA cycle-dependent growth. Mutant alleles are listed in rows and traits in columns. Cells of TF mutant alleles in heat map are colored if the AGI or AGI:TCA Metabolite linear model terms for each trait are statistically significant (P < 0.05, two-way ANOVA). Mutant alleles are hierarchically clustered using Euclidean distance. (D-F) Hypocotyl length, root length, and root to total length is dependent on TF and exogenous TCA cycle metabolites. Radar plots presents mutant phenotypes relative to Col-0 (black).

Transcription factor – TCA cycle target expression correlation analyses suggested that TCA cycle transcriptional regulation is highly context-dependent. To assess the functional contribution of select transcription factors from this sub-network to the TCA cycle and context-dependent plant growth, we obtained 31 insertional mutant alleles of 17 TFs (Key Resources). The expression of these 17 TFs was highly correlated with their TCA cycle gene targets in the salt and osmotic stress datasets, and were associated with primary metabolic GOs more than expected by chance (Table S9). To first assess their contribution to the TCA cycle, we measured hypocotyl and root length and the ratio of root length to the total seedling length (sum of hypocotyl and root lengths) of the mutant alleles. We further queried the dependence of these phenotypes on a particular stage of the TCA cycle by externally supplying TCA cycle intermediates - pyruvate, oxaloacetic acid (OAA), citrate, alpha-ketoglutarate, succinate, fumarate, and malate. We also included acetate to capture the non-cyclic pathway of the TCA cycle and sucrose as a representative respiration-dependent carbon source (Eastmond et al., 2000). In wild-type seedlings, pyruvate, OAA, citrate, alpha-ketoglutarate, fumarate, acetate and sucrose decrease hypocotyl length; while alpha-ketoglutarate, succinate, fumarate, malate and sucrose increase root length (Figure S4A and B). The ratio of root to total seedling length of wild-type Arabidopsis responded similarly to root lengths (Figure S4C). These observations support our hypotheses that TCA cycle activity and its intermediates have distinct functions in different organs (the root and hypocotyl).

TF-dependent changes and TF by TCA cycle intermediate-dependent changes in the three traits were tested using a two-way ANOVA (Figure 5B, Table S10). Of the seventeen tested TF and their corresponding mutant alleles, twelve TFs have significant genotype-dependent hypocotyl growth effects. Six TFs have significant genotype-dependent effects on root length and seven TFs have significant genotype-dependent effect on the ratio of root to total seedling length. *HD2A* and *AtHB34* are the two TFs have significant genotype-dependent effects across the three measured traits. In the majority of the TF mutant alleles tested, hypocotyl lengths decreased across the TCA intermediate conditions, indicating that the majority of these TFs promote hypocotyl length. The direction of the effect on root length and the ratio of root to total seedling length was more varied amongst the TF mutant alleles across conditions (Figure S4D), indicating the responses to the TCA intermediates in the roots and the ratio of root to total lengths are conditional to the specific TF. Over one third of the TFs have significant effects conditional on TCA intermediate supplementation (Figure 5B). Two transcription factors showed significant genotype by TCA intermediate effects in hypocotyl length - *GATA12* and *WRI3* (Figure 5C). Six of the seventeen TFs tested have significant genotype by TCA metabolite effects on root length and root to total length in the mutant allele seedlings, with five TFs common between the two traits (Figure 5B-F, Table S10). In summary, all seventeen TFs tested have an effect on plant growth in the dark and TCA-cycle dependent nutrition. Moreover, there was a near absence of TFs with significant TCA metabolite interaction effects that overlap between the hypocotyl and the root. This further supports our hypothesis that TCA cycle activity differs dependent on organ type and points to specific TFs potentially responsible for this conditionality.

### TFs affect salt stress responses

The above phenotypic analyses establishes that the TCA cycle TF sub-network affects TCA cycle-linked growth and development. We next tested our hypothesis that these transcription factors also organize TCA cycle function in response to the environment by focusing on salt stress. Salt stress generates a dramatic shift in plant primary metabolism including for the TCA cycle (Sanchez et al., 2007). Saline conditions are a major threat to agriculture and we analyzed physiologically relevant traits throughout the plant life cycle. All of the seventeen transcription factors are highly correlated with their TCA cycle target genes in salt stress conditions (Table S8). We therefore evaluated the TF mutant alleles’ response to salt stress. At seven days of age, plants were watered with either water or water with 50 mM NaCl. This concentration of salt is the minimum concentration necessary to induce a significant growth difference in wild-type Arabidopsis Col-0 seedlings (Julkowska et al., 2014). Rosette area, growth rate, dry shoot biomass, flowering time, seed yield and the natural abundance of ^13^C, ^15^N in seeds as well as their ratio (^13^C:^15^N) were measured and provide a comprehensive overview of the agriculturally-relevant consequences of TCA cycle disruption over the continuum of plant growth in response to salt stress. Transcription factor-dependent and transcription factor by salt treatment-dependent responses were evaluated using a two-way ANOVA (Table S11 and S12).

As anticipated, salt stress significantly reduced growth rate and dry shoot biomass in wild-type Col-0 plants (Figure S5A-F). Mutations in fifteen of the TCA cycle-linked TFs led to small but significant effects on shoot biomass, flowering time, growth rate and seed yield in both control and salt conditions relative to Col-0 (Figure 6A, Figure S5G). These included both positive and negative effects on rosette areas of TF mutants. Six of the TFs tested affected growth in a salt-dependent manner, including mutants of *BASIC PENTACYSTEINE 4*, *COLD SHOCK DOMAIN PROTEIN 4*, *AT2G48100*, and *CHA19* for rosette area, *LBD16* in dry shoot biomass, and *WRI3* and *AT2G48100* in seed yield (Figure 6B and 6C, Table S11 and S12).

**Figure 6.**
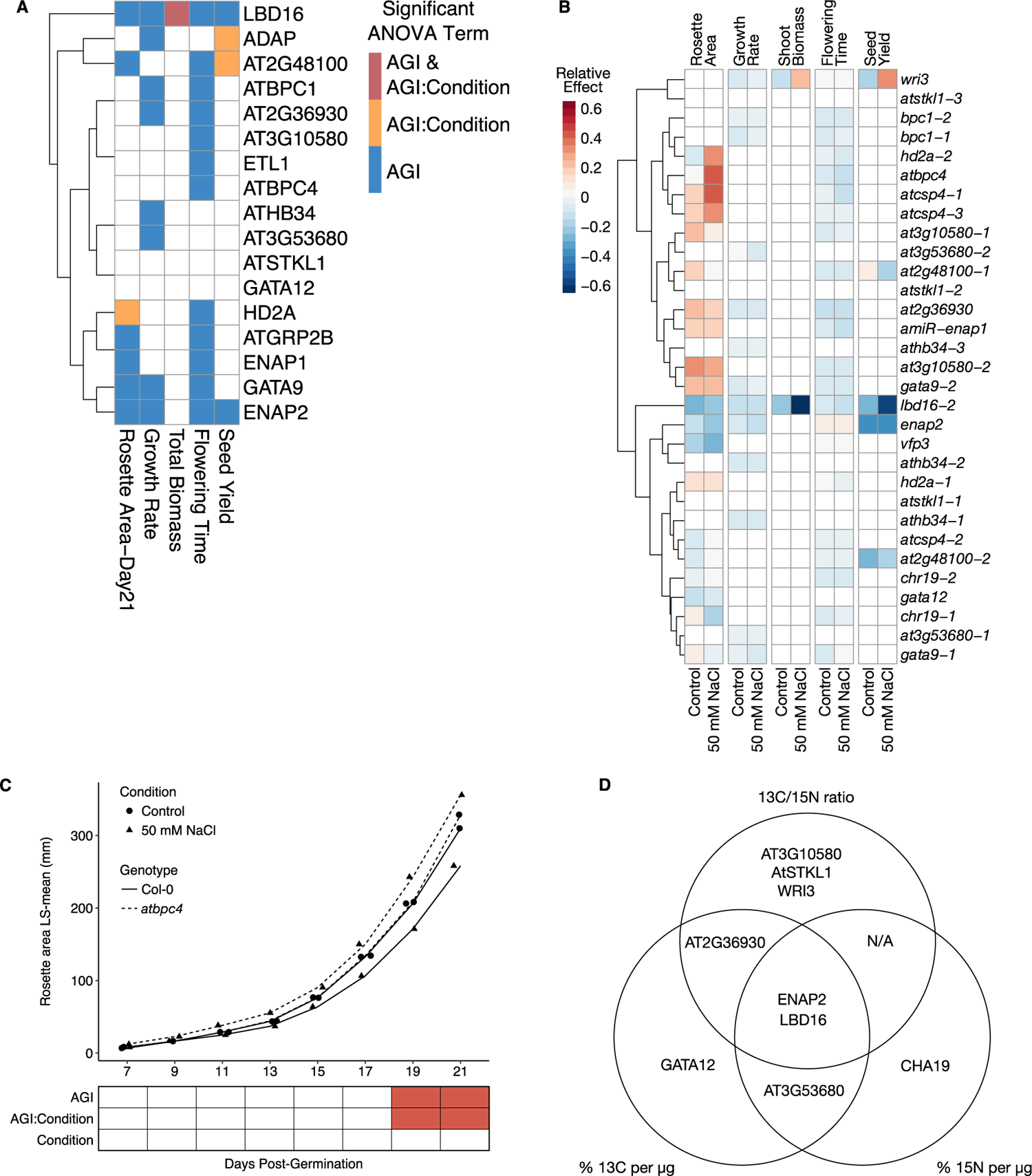
Phenotypes of mutant alleles are genotype by salt treatment-dependent. (A) Heat map summarizing the relative effect of TF mutant alleles under control and salt treatment on rosette area, growth rate, shoot biomass, flowering time and seed yield. Mutant alleles are listed in rows and traits under both control and salt treatment are in columns. Cells of TF mutant alleles in heat map are colored if the AGI or AGI:Condition term in the linear model of each trait is statistically significant (*P* < 0.05, two-way ANOVA). Mutant alleles are hierarchically clustered using Euclidean distance. (B) Rosette area of mutant allele of *BPC4* from day 7 to day 21 post-germination. Linear model for two-way ANOVA considers AGI, salt treatment, day post-germination and their interactions. The AGI and AGI:Salt Condition terms are statistically signification (*P* < 0.001, *P* < 0.01, respectively). Heat map under line plot indicates which term in linear model is statistically significant (*P* < 0.05) using two-way ANOVA for each day. (C) Venn diagram of transcription factors in which the natural abundance of ^13^C and ^15^N and the ratio of ^13^C to ^15^N that were perturbed in their respective mutant alleles. Transcription factors are listed if the AGI or AGI:Salt Treatment linear model terms are statistically significant (*P* < 0.05) as determined by a two-way ANOVA. Solid lines, Col-0; dashed line, *bpc4*; circle, control condition; triangle, 50 mM NaCl.

In addition to growth phenotypes, mutations in nine of the tested TFs led to significant changes in the abundance of ^13^C, ^15^N or the ^13^C:^15^N ratio. Five TFs have significant genotype-dependent effects on ^13^C abundance. Two TFs have significant genotype-dependent effects and two TFs have significant TF by salt treatment on ^15^N abundance. Among the TFs that affect ^13^C:^15^N ratio, two TFs have significant genotype-dependent effects and three TFs have significant effects dependent on the salt treatment. Mutations in *ENAP2* and *LBD16* generated significant differences in all three isotope abundance traits relative to Col-0 (Figure 6D, Figure S6, Table S11 and S12).

Given the influence of these TFs on both dark-mediated growth and salt stress, we investigated if the two sets of traits showed a connection as might be expected if a single metabolic pathway (the TCA cycle) underlies both context-dependent phenotypes. A positive correlation (r = 0.53, *P* = 0.03) was observed between the number of significant phenotypes observed in the dark TCA intermediate-feeding and salt stress response experiments and the number of significant C and N content phenotypes (Figure S7). In summary, our phenotypic analyses of mutant alleles of seventeen transcription factors within this TCA cycle sub-network demonstrate this regulatory module’s importance in plant growth and its response to salt stress as well as the importance of transcription factors in coordination of TCA cycle function in different organs and conditions.

## DISCUSSION

Here, we present a global map of interactions between TFs and gene promoters in central carbon and specialized metabolism in *Arabidopsis*. The yeast one hybrid methods are complementary to other high-throughput *in vitro* assays including protein binding microarrays (Weirauch et al., 2014), DAP-seq (O’Malley et al., 2016), or computational inferences (Kulkarni et al., 2018) to identify interactions between TFs and target genes. Some of these methods assay for interactions between hundreds of transcription factors - 916 TFs (Kulkarni et al., 2018) or 529 TFs (O’Malley et al., 2016). This mapped set of interactions utilize a full TF collection, which has been utilized in yeast one-hybrid assays (Bonaldi et al., 2017; Breton et al., 2016; Kang et al., 2018; Li et al., 2019), albeit with fragments of single regulatory regions. Here, we utilized an enhanced yeast one-hybrid system (Gaudinier et al., 2011) to screen for interactions with 226 promoters enabling the detection of a vastly higher proportion of transcriptional regulators of primary and secondary metabolism than these previously mentioned studies.

The resulting putative regulatory network model differs from that found in single celled organisms. In this network, most TFs bind to promoters of two or more metabolic pathways rather than being pathway-specific TFs, suggesting a model of regulation that requires TF coordination of multiple pathways. Additionally, a large collection of TFs were identified via the yeast one hybrid analyses that could potentially regulate each pathway. Together, these suggest that a pathway-specific or master-regulatory model is not likely the design principle underlying metabolism in Arabidopsis, and potentially, plants. Instead, we propose that plant metabolism is controlled via a distributed system whereby each pathway is influenced by a large collection of TFs. This collection of TFs provides the ability for multiple pathways to be coordinated as a unit and for different modules to be created. This potential was supported by the inducible transcription factor analyses whereby genes in the pentose phosphate pathway, glycine biosynthesis and cysteine biosynthesis were differentially expressed upon induction of each single TF, with enrichment for their putative regulatory targets.

An additional benefit of a distributed control system is that it allows for more fine tuning to precisely respond to either environmental or developmental cues. This potential has been previously observed in the aliphatic GLS pathway wherein TFs had tissue- and or environmentally-conditional effects (Li et al., 2014; 2018). In this study, we were able to extend this observation of developmental and environmental context-specific regulation of the TCA cycle, a central primary metabolic pathway within Arabidopsis. Genetic perturbation of seventeen TFs revealed that these factors regulate the TCA cycle and the localization of their targets suggests differential transcriptional control of TCA targets within various cellular compartments, potentially increasing the flexibility of the pathway. These findings on the TCA cycle further enhance the difference in multi-cellular metabolic regulation as the TCA cycle within the single celled *Escherichia coli* and *Saccharomyces cerevisiae* is determined by a limited number of metabolites using a few key TFs.

This resource of TF-enzyme promoter interactions in Arabidopsis has allowed us to propose a hypothesis that multi-cellular organisms rely on a distributed regulatory system for controlling metabolism rather than the previously observed master-regulatory/pathway specific system found in single celled organisms. Furthermore, it can provide an important source of regulatory modules to enable combinatorial engineering of plant metabolism across diverse metabolic pathways.

## ACKNOWLEDGMENTS

This work was supported by NSF-MCB-1330337 (to MT, BL, SMB, and DJK). S.M.B by HHMI #55108506; D.J.K. by NSF MCB #1906486 and SDA NIFA Hatch #CA-D-PLS-7033—H. M.T. by NSF GRFP and the UC Davis Department of Plant Sciences Graduate Research Assistantship.

## AUTHOR CONTRIBUTIONS

D.J.K, S.M.B, M.T. and B.L. conceived and designed the experiments. D.J.K and S.M.B. secured funding for the project. M.T. and B.L. performed the promoter and TF cloning. M.T., B.L. and X.Z. performed the Y1H experiments. M.T., B.L., X.Z., T.B., J.J.L., N.C., A.G., R.N., C.C.W. performed the phentoyping experiments. M.T., T.B. and J.J.L performed the RNA-Seq experiments. M.T. analyzed the data. MT, S.M.B and D.J.K wrote the manuscript with contributions from all authors. D.J.K and S.M.B. supervised the project.

## DECLARATION OF INTERESTS

The authors declare no competing interests.

## STAR METHODS

### EXPERIMENTAL MODEL AND SUBJECT DETAILS

#### Saccharomyces cerevisiae (Transformation)

*Saccharomyces cerevisiae* strain YM4271 (for promoters) or strain Y*α*1867 (for TFs) were grown overnight at 30°C by taking streaking from a glycerol stock onto a YPDA agar plate to make a lawn. A pea-sized glob of yeast cells was resuspended in 1 mL of liquid YPDA and ∼100 μL was added to 50 mL of liquid YPDA in an Erlenmeyer flask to a starting OD_600_ of 0.15-0.20. The yeast culture was grown in a 30°C shaking incubator at 210 rpm for two hours or until OD_600_ had reached 0.4-0.6.

#### Saccharomyces cerevisiae (Y1H)

Five mL of SC-HIS/-URA was inoculated with yeast strains of the promoters from glycerol stock and grown at 30°C and shaking at 210 rpm for 48 h. Cultures of promoter yeast strains were concentrated by pelleting down yeast cells (centrifuge 5 min at 1850 x g) and resuspending in 1-1.5 mL of liquid SC-HIS/-URA, and then 510 μL spread onto SC-HIS/-URA agar plates with 15-20 glass beads. Promoter lawn plates were incubated for 2 nights at 30°C. The TF collection was cultured in deep 96-well plates from glycerol stocks in 315 μL of liquid SC-TRP for two nights at 30°C. TF culture plates were spun down for 5 min at 1850 x g and 150 μL of liquid media was removed. The pellets were resuspended in the remaining liquid media and 50 μL of the culture transferred to 384-well plate and arrayed in duplicates, resulting in two 96-well plates combined into one 394-well plate.

#### *Arabidopsis thaliana* (Bulking and genotyping)

T-DNA insertional mutant lines were ordered from the Arabidopsis Biological Resource Center at Ohio State University (Alonso et al., 2003) or obtained from published sources (Zhang et al., 2016). To minimize for variation due to maternal conditions, all mutant lines were bulked together with wild-type except for *at2g48100-2*, *enap1* and *gata12* (Key Resources), which were bulked together with another growth of the wild-type reference. Arabidopsis plants were grown in Sunshine mix #1 in 2.23” x 1.94” x 2.23” pots. Chamber temperature was set to continuous 22°C and set to long day (16-hr light/8-hr dark) with light intensity of ∼120 umol m^-2^s^-1^ from T12 VHO fluorescent light bulb. Seeds were sieved for size between 250-300 microns and surface sterilized by soaking in 50% household bleach, 0.05% Tween 20 for 20 minutes and then rinsed with sterile water 5-7 times before stratifying in 0.1% sterile agar for 2 days in 4°C. Seeds were sown in Sunshine Mix #1 in traditional 1020 trays with 18 cell inserts (pot dimension 3.10” x 3.10” x 2.33”).

#### *Arabidopsis thaliana* (TCA metabolite feeding)

Seeds were plated onto half-strength MS agar pH 5.7 with KOH, supplemented with 1 mM TCA metabolites or 0.21 mM sucrose based on theoretical ATP yield (Rich, 2003). The concentration of sucrose was calculated to provide an equivalent theoretical yield of ATP as 1 mM of the TCA metabolites. pH of 0.25 M OAA, keto-glutaric acid, and pyruvic acid were adjusted with KOH, filter sterilized (0.22 μm) and then added to MS agar after autoclaving to a final concentration of 1 mM. Plates were then wrapped in double layers of aluminum foil and grown standing vertically for 5 days in the dark in ambient room temperature (21-23C). Germination was determined by unwrapping additional plates each day post-plating.

#### Arabidopsis thaliana (Salt response)

Seeds were sieved for size between 250-300 microns and surface sterilized by soaking in 50% household bleach, 0.05% Tween 20 for 20 minutes and then rinsed with sterile water 5-7 times before stratifying in 0.1% sterile agar for 2 days in 4°C. Seeds were sown in Sunshine Mix #1 in traditional 1020 trays with 18 cell inserts (pot dimension 3.10” x 3.10” x 2.33”) and grown in chamber with temperature set to continuous 22°C and set to long day (16-hr light/8-hr dark) with light intensity of ∼120 umol m^-2^s^-1^ from T12 VHO fluorescent light bulb. After germination, each pot was thinned to one seedling per pot. When seedlings were 7 days old, flats were watered every 5-7 days with either deionized water (control) or 50 mM NaCl (liquid; treatment) to maintain soil moisture.

#### Arabidopsis thaliana (Transformation)

Ten to twelve *Arabidopsis thaliana* Col-0 seeds were sown in Sunshine Mix #1 in 3.5” x 3.5” x 5” pots. Pots were incubated at 4°C for 3 days before transferring into growth chamber set to continuous 22°C and set to long day (16-hr light/8-hr dark) with light intensity of ∼120 umol m^-2^s^-1^ from T12 VHO fluorescent light bulb. Five days after seed germination, each pot was thinned to five seedlings per pot and grown to four- to five-weeks for Agrobacterium transformation.

#### Arabidopsis thaliana (RNA-Seq)

T3 seeds from four independent lines were selected by presence of RFP in the seed. Approximately 200-300 seeds of each line were surface sterilized with 50% bleach for 15 minutes, rinsed 5-7 times with sterile distilled water, and imbibed in the dark at 4°C for two days. Seeds were then sowed onto nylon mesh on Petri plates containing half-strength MS agar plates pH 5.7 with KOH. Plates were double wrapped in foil and placed vertically in a dark chamber at ambient room temperature (20-22 C).

## METHOD DETAILS

### Network visualization

All networks are visualized using Cytoscape v3.7.1 (Shannon et al., 2003).

### Bait Promoter Cloning

PCR primers were designed to amplify promoter regions of 2000 bp in size or to the next gene upstream from the predicted translational start site of each TCA cycle gene (Table S1). Promoter regions were amplified from Col-0 genomic DNA or from plasmids containing 1 kb synthesized promoters (Life Technologies) using Phusion High-Fidelity Taq (NEB) and cloned into pENTR 5’ TOPO vector (Invitrogen). Promoter regions were then recombined with pMW2 and pMW3 destination vectors (Deplancke et al., 2004) in LR reactions and sequence-confirmed before transforming into the YM4271 yeast strain as described in (Gaudinier et al., 2011).

### Prey TF Cloning

cDNA of TFs in pENTR from (Pruneda-Paz et al., 2014) that were not present in the root-expressed TF collection (Gaudinier et al., 2011) were recombined with pDEST-AD-2micron destination vector in LR reactions. Prey TFs were transformed into Yα1867 yeast strain.

### Yeast One-Hybrid

All TCA cycle promoter baits were screened against a total collection of 2039 prey TF according to protocols in (Gaudinier et al., 2011; 2017).

### TF Family Enrichment

Subcellular localization of TCA targets was determined from the SUBA3 database (Tanz et al., 2013). The consensus predicted subcellular localization was used except when a gene’s subcellular localization was previously experimentally tested as per the literature. In that case, the experimentally determined subcellular localization was used. The number of interactions was obtained for each TF family targeting a TCA enzymatic step in each cellular compartment from the yeast one-hybrid interaction data. TF family enrichment was tested using the Fisher’s exact test with Holm–Bonferroni to adjust for multiple testing in R. The number of TFs in each cellular compartment in each family was tested against the number of TFs in the yeast one-hybrid assay.

### Correlation analysis

The yeast one-hybrid data generated a network of TF-TCA target gene interactions, where TF or target genes were nodes and interactions were edges. Using this network structure, Pearson correlation coefficients were calculated for each TF-TCA gene interaction in R using the cor function with gene expression values from publicly available microarray datasets (Key Resources). These datasets capture a range of experimental and biological perturbations and thus allow for the capture of conditional transcriptional correlations. We downloaded available .CEL files from the experiments only when the genotype corresponded to *Arabidopsis thaliana* accession Col-0. This was to exclude the complications of polygenic natural variation between different accessions and to focus on perturbations in a single genotype. Using the .CEL files, the microarray datasets were normalized using the rma function from the affy package (Gautier et al., 2004) in Bioconductor.

We parsed these datasets to represent specific aspects of developmental biology or stress response. The first was the Arabidopsis development (Schmid et al., 2005) and root development expression atlases (Birnbaum et al., 2003; Brady et al., 2007; Lee et al., 2006; Levesque et al., 2006) that encompass numerous cell-, tissue- and organ-types. We added the pollen development expression atlas (Honys and Twell, 2004; Qin et al., 2009) because of the coordinated transcriptional changes with strong temporal regulation observed in pollen metabolism, across pollen maturation and pollen germination. In addition to development, we included a large salt and osmotic stress (Kilian et al., 2007) expression atlas because these stresses modulates photosynthesis and cellular respiration associated with the TCA cycle.

### Gene Ontology Enrichment Analysis

AGI loci identifiers (see Table S6) were uploaded to the GO Term Enrichment tool powered by PANTHER on www.arabidopsis.org.

### TCA Metabolite Feeding Assay

For each experiment, 30 replicates of Col-0 and 10 mutants were plated in a random block design. TF mutant lines were divided into two blocks. Each plate (block) had 6 wild type seedlings and one seedling per mutant lines for 15 or 16 TF mutant genotypes plated across 3 rows. The entire experiment was repeated twice, for a total of 20 biological replicates maximum per TF mutant lines and 60 biological replicates maximum of Col-0 per TCA metabolite. Backlit images of plates on a light table were acquired using a Canon EOS T3i Rebel dSLR camera fixed on a camera stand. Roots and hypocotyls were traced manually using a Wacom Intuos drawing tablet in ImageJ.

### Salt response assay

Mutant seedlings were grown in a random block design. Each block consists of 10 to 11 mutant TF lines plus Col-0, for a total of 3 blocks. Each experiment consists of 5 biological replicates per TF mutant lines and the entire experiment was repeated twice for a total of 10 biological replicates maximum per TF mutant. The concentration of salt selected was the minimum concentration necessary to induce a significant growth difference in wild-type Arabidopsis Col-0 seedlings (Julkowska et al., 2014). A copy stand and Canon T3 dSLR were used to take aerial images of each flat every other day starting at 7 days post-germination until 21 days post-germination. Each image had a measurement marker to standardize size images between pictures. Image analysis to obtain rosette area was conducted in ImageJ using the Analyze Particle function after selecting Hue (42-166), Saturation (28-255), Brightness (80-255) under Adjust > Color Threshold. Dry shoot biomass was determined from fully mature shoots from 3 to 5 individuals of each genotype at the end of the first salt stress experiment. Shoots were dried in 60°C oven overnight and then weighed on an analytical balance.

### Natural ^13^C and ^15^N abundance profiling

For Carbon-13 and Nitrogen-15 profiling, mature seeds from 3 to 5 individuals of each genotype were sieved and cleaned to remove any plant material and then dried in a 60 C oven overnight. The seeds were obtained from the control and salt conditions from the first salt stress experiment. Seeds were allowed to acclimate to ambient room conditions before two to three mg of seeds were submitted to the Stable Isotope Facilities at UC Davis for determining natural abundance of Carbon-13 and Nitrogen-15.

### Statistical analysis for T-DNA insertional mutant phenotype assays

Replication numbers for all experiments were designed to provide significant power for moderate effect sizes and modest power for small effect sizes using a presumed broad-sense heritability of about 15% based on previous experience with these traits. All statistical analyses were conducted in R version 3.4.0. The TFs controlling root length, hypocotyl lengths, root to total seedling lengths, rosette area, vegetative growth rate, dry shoot biomass, flowering time and seed yield were tested by ANOVA using a general linear regression model in R with the package lmerTest. The following generalized nested linear mixed model was used: Trait = AGI + AGI:Allele + Treatment + AGI:Treatment + AGI:Allele:Treatment. In the case of rosette area, day was included as a factor: Trait = AGI + AGI:Allele + Treatment + Day + AGI:Day + AGI:Treatment + Treatment:Day + AGI:Allele:Treatment + AGI:Treatment:Day. The random effect terms for the dark feeding experiment are Block nested in Experiment and Row on Plate nested in Plate. The random effect terms for the salt response experiment are Flat Condition nested in Shelf nested in Experiment. Random-effect terms in the models were computed using ranova function from the lmerTest package. To allow for AGI:Allele term in the generalized linear model, growth traits (hypocotyl lengths, root lengths and rosette areas) of the two alleles that were bulked at a different time were normalized by adding the difference between the two wild types to the raw values. Estimated marginal means (EMMs/Least square means) and post hoc comparisons between mutant alleles and Col-0 conditioned on Treatment were calculated using the emmeans (Lenth, 2016) package in R. P-values from post hoc tests were adjusted for multiple comparisons using the Holm method in R. Relative effect is expressed as EMM_Mutant_ – EMM_WT_/EMM_WT_.

### Cloning dexamethasone-inducible overexpression TF lines

Coding sequences of *CHA19*, *ENAP1*, *LBD16* and *WRI3* in pENTR from the Arabdiopsis TF ORFeome collection (Pruneda-Paz et al., 2014) (CD4-88) were recombined into a modified pFAST-R05 plasmid with the dexamethasome-inducible GR construct from pBEACONRFP-GR plasmid using LR reactions. To generate the modified vector, pFAST-R05 (Shimada et al., 2010) was digested with SbfI and ApaI to serve as the backbone of the modified plasmid, containing the LB, RB and pOLE1:OLE1-RFP, a seed-selectable RFP marker. The dexamethasone-inducible overexpression cassette from pBEACONRFP-GR (Bargmann et al., 2013) was cloned using Phusion High Fidelity Taq (Forward primer: GACTAGAGCCAAGCTGATCTCC; Reverse primer: CGACGTCGCATGCCTGCAGG), sequenced verified and then recombined into the digested pFAST-R05 backbone using Gibson assembly (NEB E2611S) (Forward primer: CCTGCAGGCATGCGACGTCGTCAAGCTTAGCTTGAGCTTGGATCA; Reverse primer: CAAGCTCAAGCTAAGCTTGACGACGTCGCATGCCTGCAGG). Recombined TFs in the modified pFAST-R05 with the dexamethasone-inducible GR system were sequence-confirmed and then transformed into *Agrobacterium tumefaciens* strain EHA105 using the calcium chloride freeze-thaw method (Holster et al., 1978). Transformed Agrobacterium were selected on LB agar plates containing spectinomycin and rifampin and then confirmed by genotyping polymerase chain. Four- to five-week old, flowering *Arabidopsis thaliana* Col-0 were transformed using floral dip method (Clough and Bent, 1998).Transformants were selected by presence of RFP in the seed coat using a 532 nm green laser and red filter, and confirmed by genotyping PCR. Transformants were bulked together to T3 under the same conditions as the T-DNA insertional mutant lines.

### GR-induction and RNA-Seq library construction

At day 5, seedlings on nylon mesh were transferred onto mock plates or half-strength MS agar plates containing 10 μM dexamethasone (dex) in the dark, assisted with a green headlamp. After 24 hours of induction, whole seedlings were collected, flash frozen in liquid nitrogen and stored in −80°C until RNA-Seq library constructions. Direct mRNA isolation was performed using biotinylated polyT oligonucleotide and Streptavidin-coated magnetic beads following protocol from (Townsley et al., 2015). Non-strand specific RNA-Seq libraries were prepared according to the protocol for high-throughput RNA-Seq library preparation method (Kumar et al., 2012). A total of 64 RNA-Seq libraries were made, representing two bioreps for the two treatments across four independent dex-controlled GR-TF mutant lines (Key Resources). Libraries were sequenced twice on the Illumina HiSeq 4000 system in SR100 mode at University of California, Davis DNA Technologies Core.

### RNA-Seq Analysis

FastQ file processing was performed on the University of California, Davis high performance bioinformatics cluster. FastQ files were quality checked using FastQC v.0.11.7 (https://www.bioinformatics.babraham.ac.uk/projects/fastqc/), adapter and poly-A trimmed using TrimGalore v0.6.0 using default settings (http://www.bioinformatics.babraham.ac.uk/projects/trim_galore/) and then re-assess for quality using FastQC. Gene counts were obtained by mapping trimmed reads to the *Arabidopsis thaliana* genome (TAIR10) using STAR aligner v2.7.0f (Dobin et al., 2013). Assessing that the read count and quality were similar across the two sequencing run, gene counts were summed up. Statistical analysis was performed in R 3.6.0 using the octopus pipeline established in (Zhang et al., 2017) (https://github.com/WeiZhang317/octopus). Briefly, trimmed mean of M value normalization was performed using the calcNormFactors function from edgeR (Robinson et al., 2010) version 3.26.6. For each TF, we ran the following negative binomial generalized linear model using the glm.nb function from MASS package to test for genes that were significantly influenced by the translocation of the TF into the nucleus: *Y = T_I_ + A_L_ + T_I_ × A_L_* where the main effects T and A are denoted as dexamethasone treatment and allele of TF, respectively. False discovery was corrected using Benjamini Hochberg using p.adjust function. Gene ontology analysis was conducted on pantherdb.org by submitting the list of AGIs of genes significantly differentially expressed due to translocation of the GR-TFs chimeric protein into the nucleus. The complete biological process GO annotation dataset for *Arabidopsis thaliana* was used for testing statistical overrepresentation. Enrichment analysis was performed in R using the fisher.test function.

---

**Figure S1.**
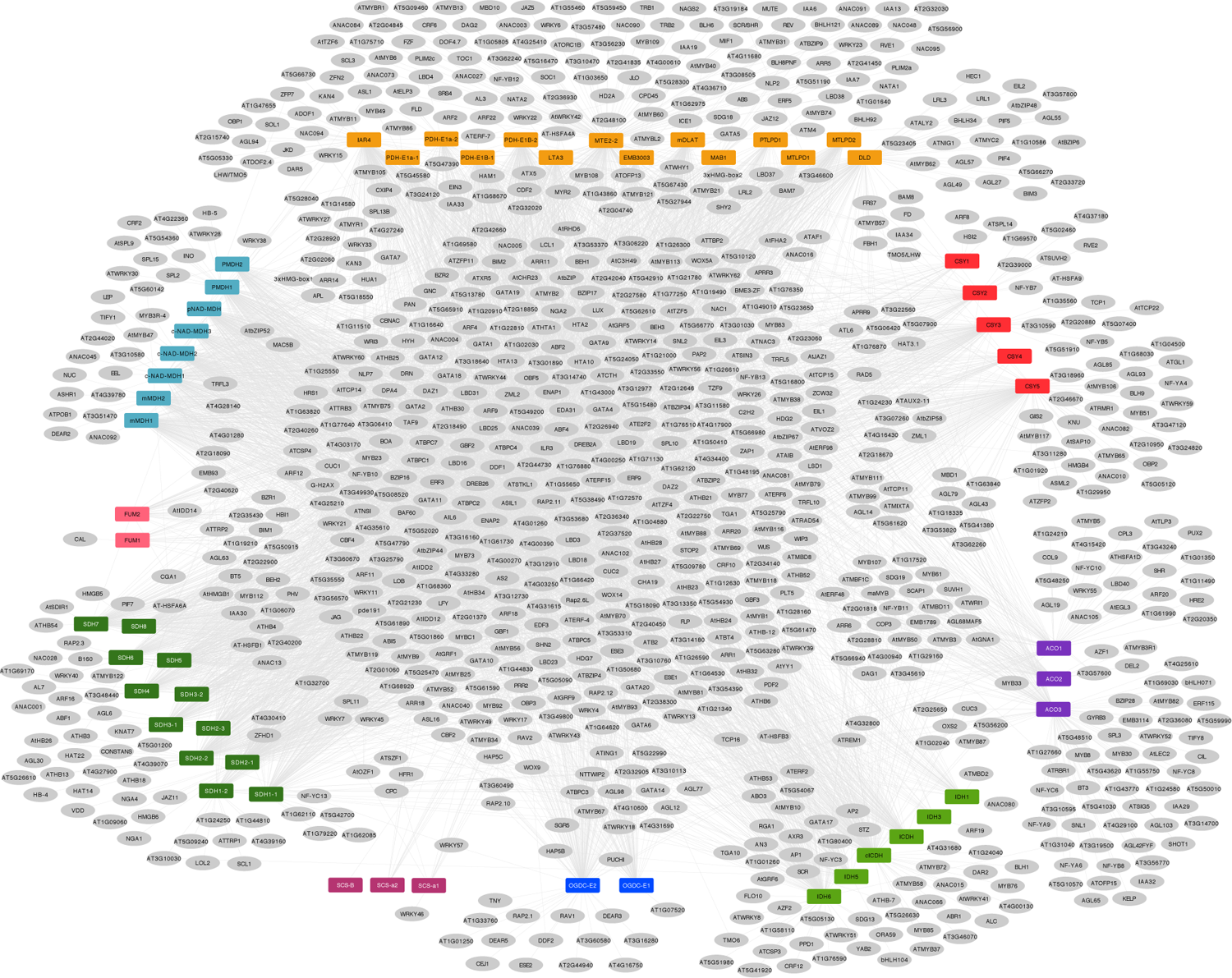
TF-TCA target gene interaction network. Yeast one-hybrid assays revealed a large and combinatorial network of TF-TCA target gene interactions. Colored rectangles, promoters; grey oval, transcription factor; grey edge, interaction. Orange PDC, pyruvate dehydrogenase; red CSY, citrate synthase; purple ACO, aconitase; light green IDH, isocitrate dehydrogenase; blue OGD, oxoglutarate dehydrogenase; light purple SCL, succinyl CoA ligase; green SDH, succinate dehydrogenase; pink FUM, fumarase; light blue MDH, malate dehydrogenase.

**Figure S2.**
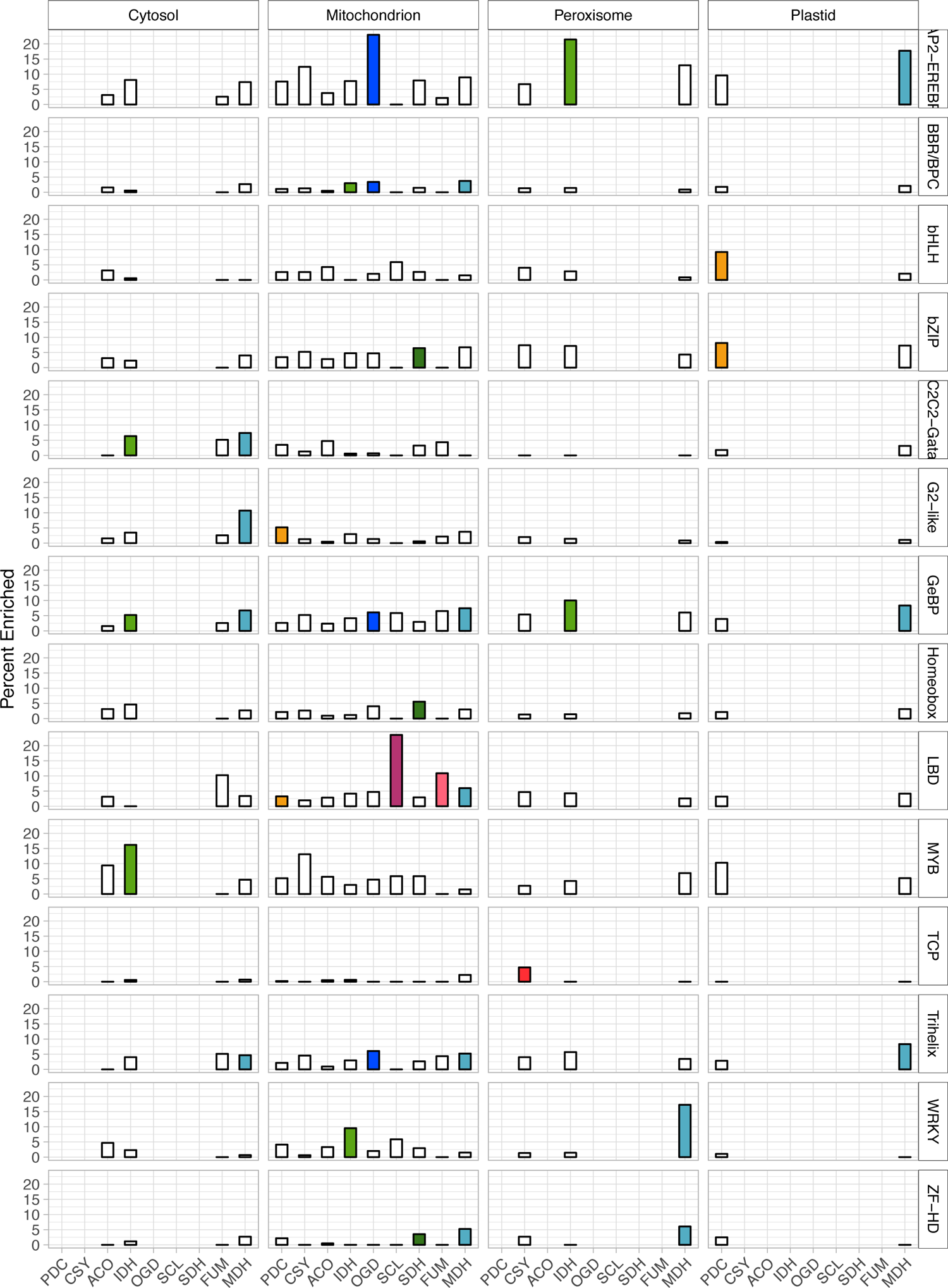
TF families are enriched for TCA cycle gene targets in specific cellular compartments. Bar graphs display the percent of TFs enriched for binding to promoters of TCA cycle enzyme in the cytosol, mitochondrion, peroxisome and plastid. Bar was colored if the TF family targeting TCA cycle enzyme in the cellular compartment was significant (adjusted *P* < 0.05, Fisher’s Exact Test, Table S7). Colors of bars correspond to the TCA enzyme in Figures 4 and S1.

**Figure S3.**
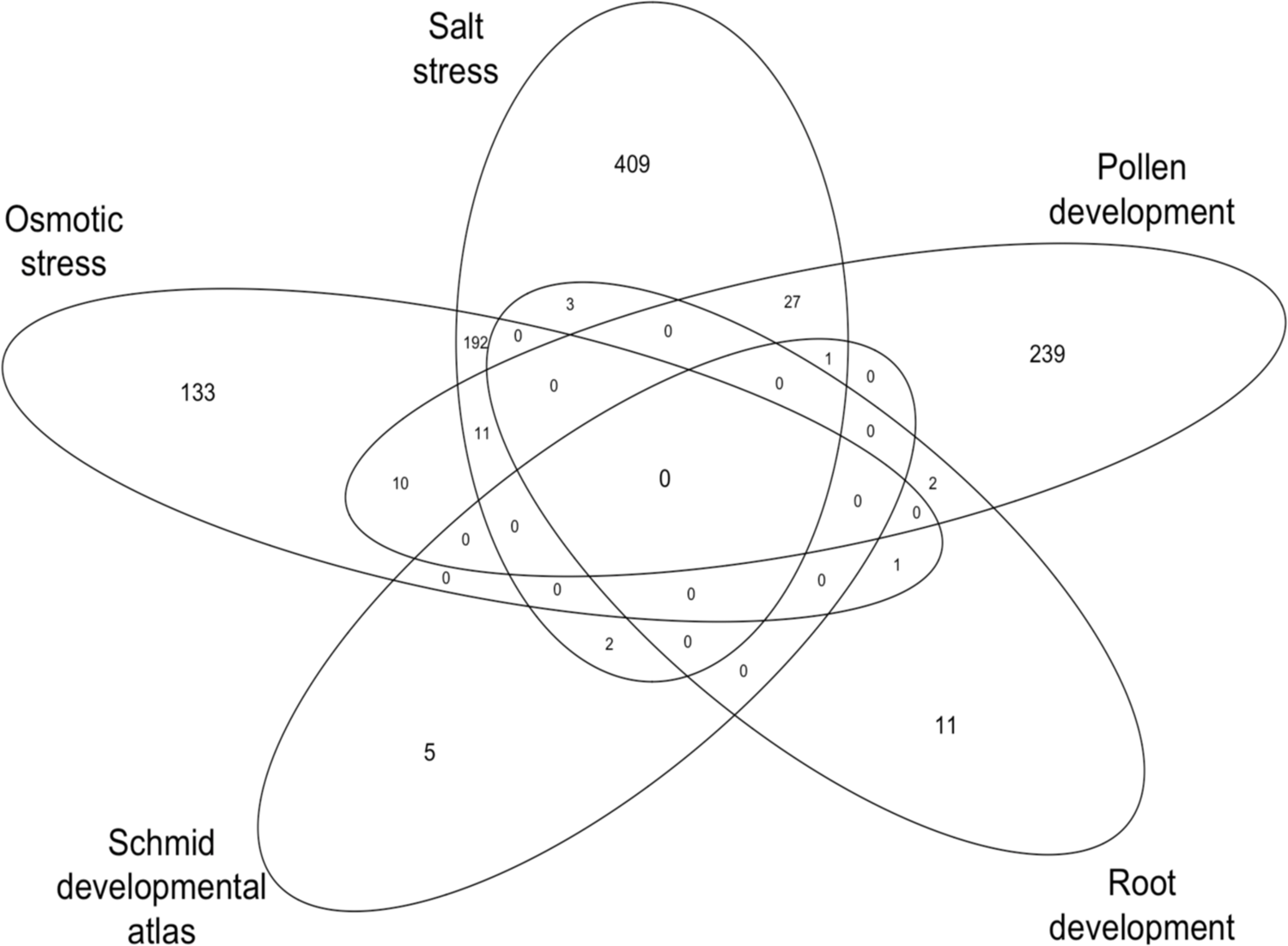
Highly correlated TF-TCA target gene interactions shared between microarray data sets. Numbers in Venn diagram represent TF-TCA target gene interactions with the absolute value of Pearson correlation coefficient ≥ 0.8 (Table S8).

**Figure S4.**
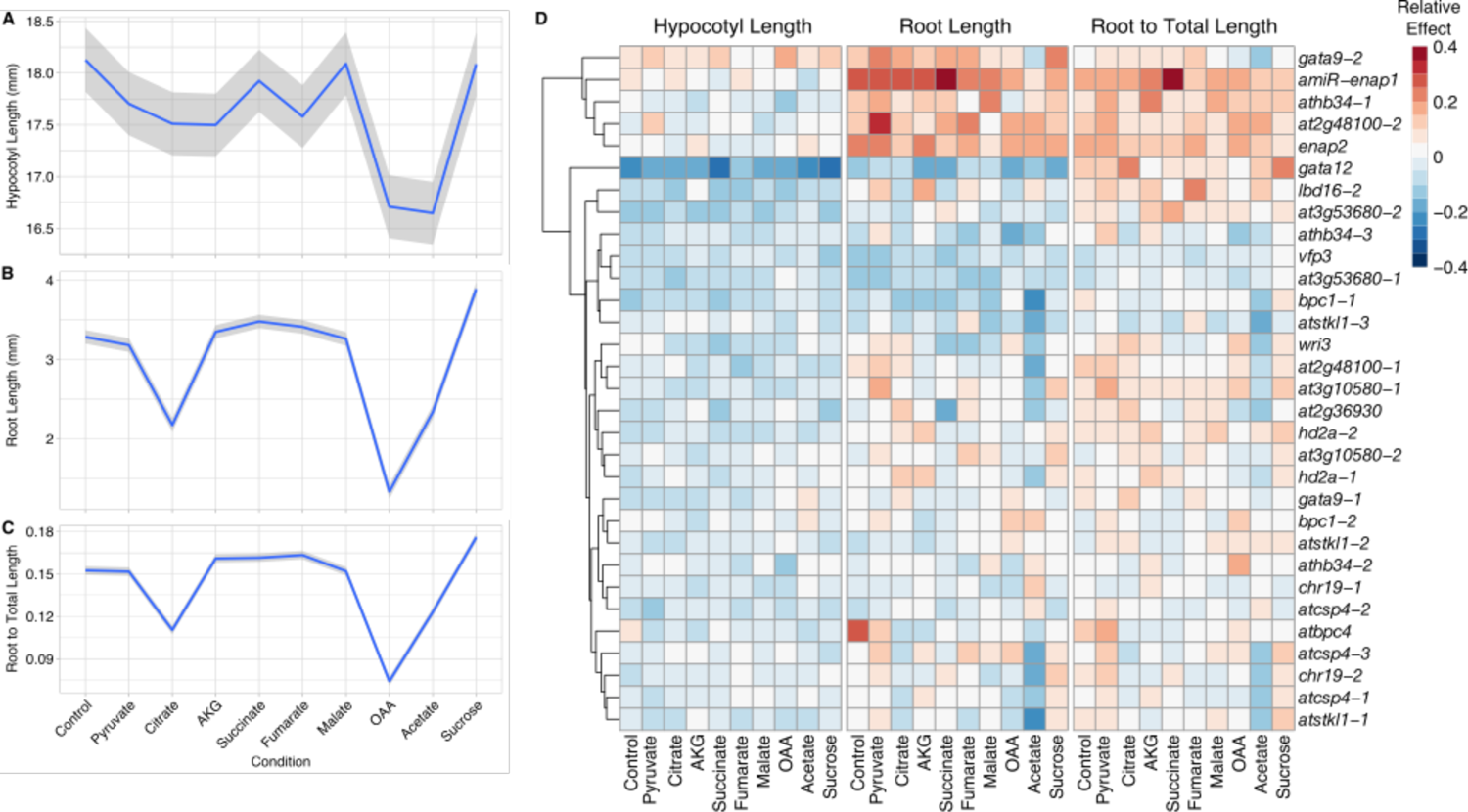
Wild-type Col-0 and TF mutant responses to TCA metabolites. (A, B and C) Hypocotyl length, root length and the ratio of root to total length of *Arabidopsis thaliana* Col-0 seedlings grown on control or TCA metabolites. Data shown is mean (blue) ± SE (grey). (D) Heat map summarizing the relative effect of TF mutant alleles on hypocotyl length (left), root length (center) and ratio of root to total length (right) on control or TCA metabolites-supplemented media. Mutant alleles are listed in rows.

**Figure S5.**
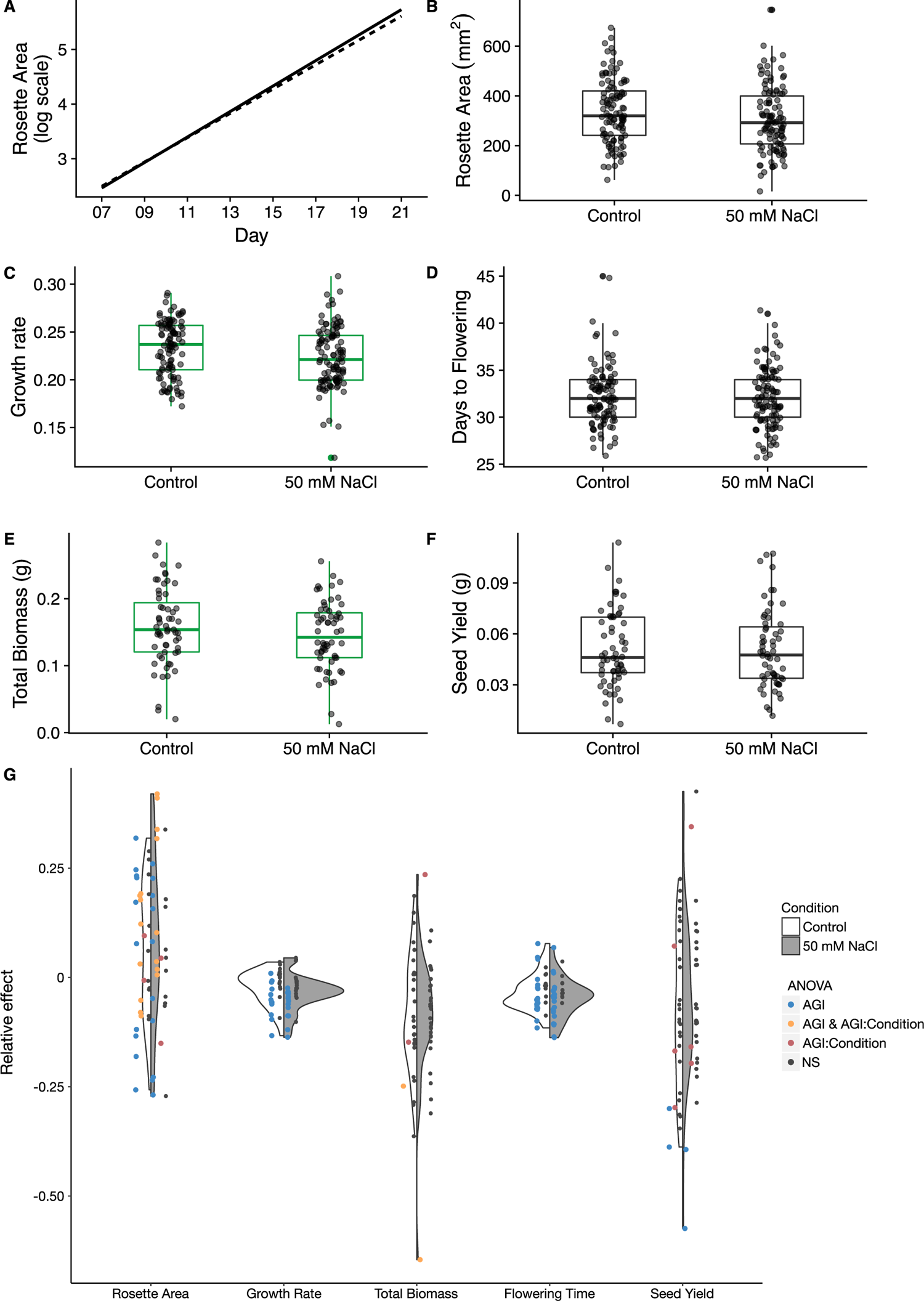
Wild-type Col-0 and TF mutant responses to salt treatment. (A-F) Rosette area, growth rate, days to flowering, dry shoot biomass and seed yield of wild-type Col-0 under control and salt condition. Solid line, control and dashed line, 50 mM NaCl. Green-colored boxplots indicate significant differences between treatments (*P* < 0.05, Student’s t-test). (G) Distribution of relative effects of TF mutant alleles under control and salt stress conditions. Dots represent TF mutant alleles. TF mutant alleles are colored if AGI (blue), AGI:Condition (red) and AGI and AGI:Condition (orange) linear model terms are significant (*P*<0.05, two-way ANOVA). White, control condition; gray, salt stress condition. Boxplots mark the interquartile range, from the 25th to the 75th percentile, and are centered at the median. Whiskers extend to 1.5*interquartile range below the lower quartile and above the upper quartile. Grey dots represent individual measurements.

**Figure S6.**
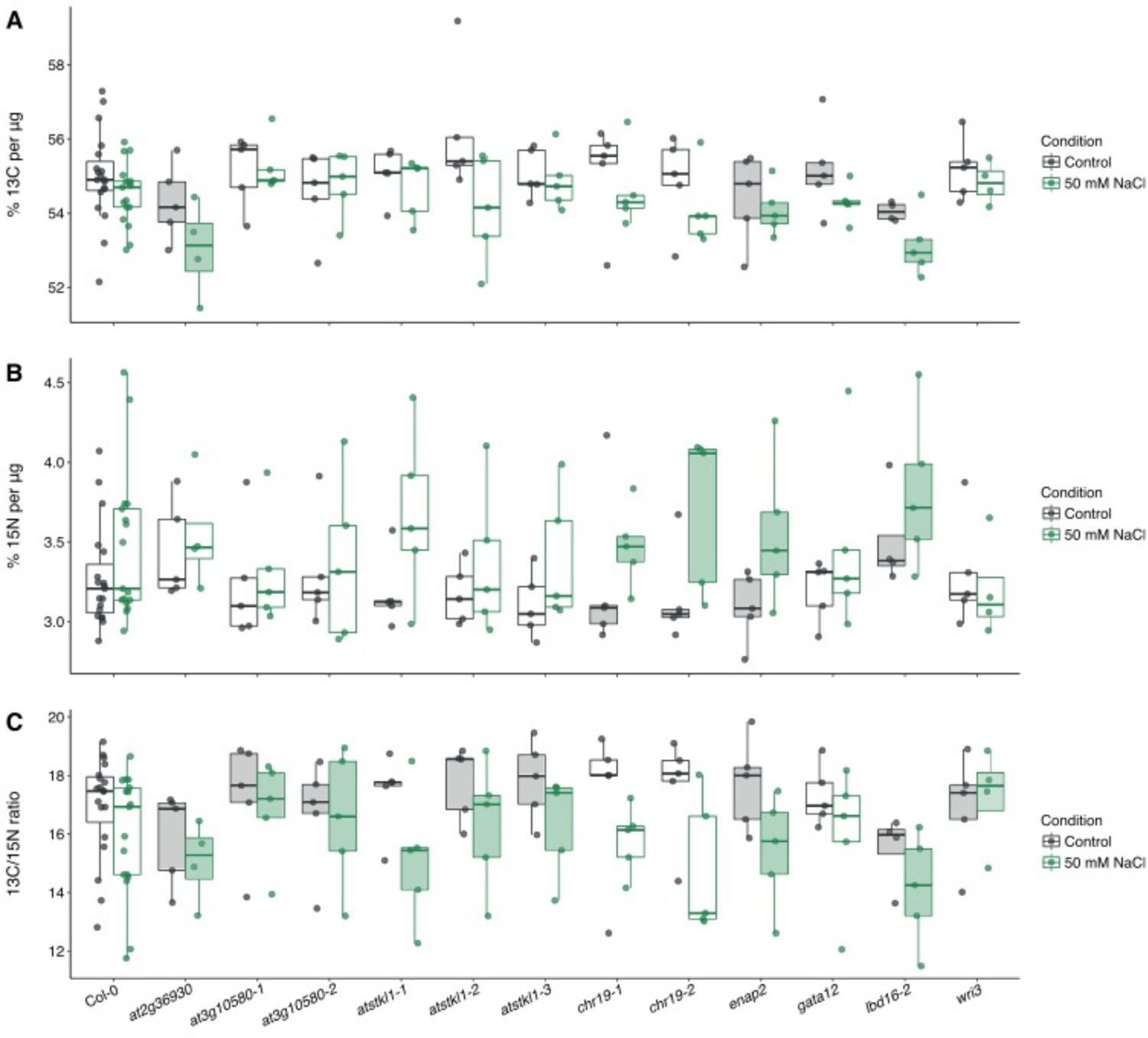
^13^Carbon, ^15^nitrogen and carbon:nitrogen ratio of TF mutant alleles. (A, B and C) Percentages of ^13^C, ^15^N and ratio of ^13^C:^15^N of seeds of individuals grown in control and salt stress conditions. Box plots are shaded in if AGI or AGI:Condition linear model terms were significant (*P* < 0.05, two-way ANOVA). Boxplots mark the interquartile range, from the 25th to the 75th percentile, and are centered at the median. Whiskers extend to 1.5*interquartile range below the lower quartile and above the upper quartile. Individual measurements are plotted as dots.

**Figure S7.**
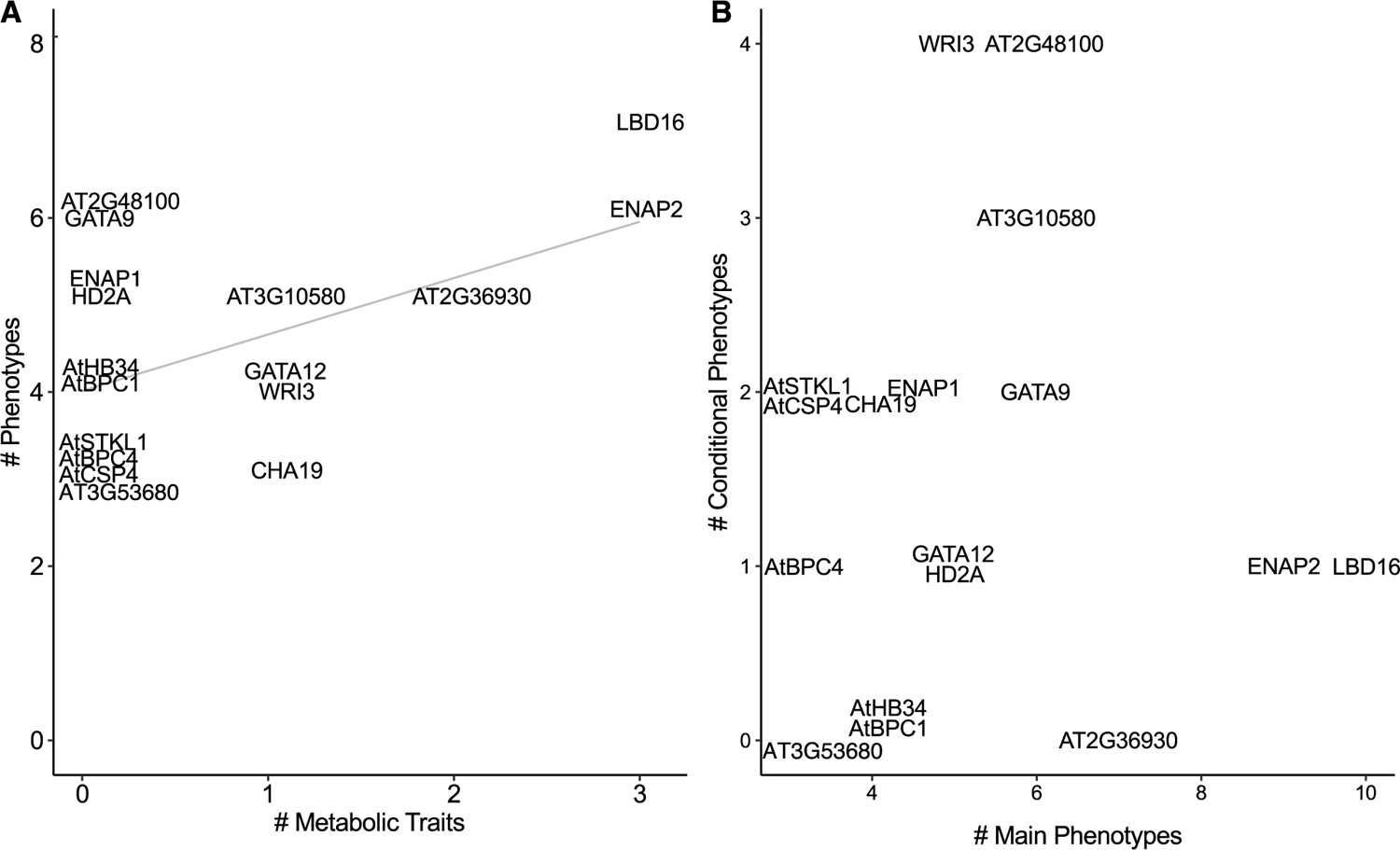
Summary of TF mutant growth traits and metabolic phenotypes. (A) A positive correlation is observed between the number of significant growth phenotypes from the TCA metabolite feeding and salt stress experiments and the number of significant metabolic phenotypes of seeds from the salt stress experiment (r = 0.53, R^2^ = 0.28, *P* = 0.03). (B) No correlation is observed between the total number of TFs with significant main effects and the total number of TFs with significant conditional (interaction) effects (r = 0.034, R^2^ = 0.0011, *P* = 0.90).

## SUPPLEMENTAL INFORMATION

### SUPPLEMENTAL TABLES

**Table S1.** Locus, length, pathway annotation and oligonucleotides for PCR amplification of promoters in this study.

**Tables S2.** Locus and source of TFs in this study.

**Table S3.** Y1H network of interactions between TFs and promoters of CCP and aliphatic GSL genes.

**Table S4.** Pairwise comparison of metabolic pathways and the number of TFs identified.

**Table S5.** GO analysis of TFs identified for each metabolic pathway.

**Table S6.** ANOVA tables and expression values from RNA-Seq analysis of GR-TFs.

**Table S7.** TF family enrichment of the TCA cycle Y1H network.

**Table S8.** TF-TCA target correlation analysis for salt stress, osmotic stress, root development, development atlas and pollen development.

**Table S9.** Gene ontology enrichment of tested TFs.

**Table S10.** Dark TCA metabolite feeding experiment results. ANOVA tables for testing hypocotyl length, root length and root to total length in the dark TCA metabolite feeding assay (A). Estimated marginal means, standard error and 95% confidence levels of hypocotyl length (B), root length (C), ratio of root to total length (D) of TF mutant lines in the dark TCA metabolite feeding assay.

**Table S11.** ANOVA tables for rosette area (A), growth rate (B), dry shoot biomass (C), days to flowering (D), seed yield (E), ^13^C ratio (F), ^15^N ratio (G), and ^13^C/^15^N ratio (H).

**Table S12.** Estimated marginal means, standard error and 95% confidence levels of rosette area (A), growth rate (B), dry shoot biomass (C), days to flowering (D), seed yield (E), ^13^C ratio (F), ^15^N ratio (G), and ^13^C/^15^N ratio (H).

## Notes

### Competing Interest Statement

The authors have declared no competing interest.

